# Health and population effects of rare gene knockouts in adult humans with related parents

**DOI:** 10.1101/031641

**Authors:** V. Narasimhan, K.A. Hunt, D. Mason, C.L. Baker, K.J. Karczewski, M.R. Barnes, A.H. Barnett, C. Bates, S. Bellary, N.A. Bockett, K. Giorda, C.J. Griffiths, H. Hemingway, Z. Jia, M.A. Kelly, H.A. Khawaja, Monkol Lek, S. McCarthy, R. McEachan, K. Paigen, C. Parisinos, E. Sheridan, Laura Southgate, L. Tee, M. Thomas, Y. Xue, M. Schnall-Levin, P.M. Petkov, C. Tyler-Smith, E.R. Maher, R.C. Trembath, D.G. MacArthur, J. Wright, R. Durbin, D.A. van Heel

## Abstract

Complete gene knockouts are highly informative about gene function. We exome sequenced 3,222 British Pakistani-heritage adults with high parental relatedness, discovering 1,111 rare-variant homozygous likely loss of function (rhLOF) genotypes predicted to disrupt (knockout) 781 genes. Based on depletion of rhLOF genotypes, we estimate that 13.6% of knockouts are incompatible with adult life, finding on average 1.6 heterozygous recessive lethal LOF variants per adult. Linking to lifelong health records, we observed no association of rhLOF genotypes with prescription- or doctor-consultation rate, and no disease-related phenotypes in 33 of 42 individuals with rhLOF genotypes in recessive Mendelian disease genes. Phased genome sequencing of a healthy *PRDM9* knockout mother, her child and controls, showed meiotic recombination sites localised away from *PRDM9*-dependent hotspots, demonstrating *PRDM9* redundancy in humans.

## Main Text

The study of gene function by correlating genotype with phenotype has provided profound biological insights for several decades. In particular, complete gene knockouts, typically caused by homozygous loss of function (LOF) genotypes, have been used to identify the function of many genes, predominantly through studies in model organisms and of severe Mendelian-inherited diseases in humans. However, information on the consequences of knocking out most genes in humans is still missing. Naturally occurring complete gene knockouts, whilst exceptional in outbred populations, offer the opportunity to study directly the lifelong effects of systemic gene inactivation in a living human. They provide a key entry point into human biology that can then be tested in other systems, and can lead to direct validation of specific biological hypotheses. The first large survey of LOF variants in adult humans demonstrated ~100 predicted LOF genotypes per individual, describing around ~20 genes carrying homozygous predicted LOF alleles and hence likely completely inactivated (*1*). As expected almost all these homozygous genotypes were common (allele frequency) variants, and were concentrated in genes likely to have weak or neutral effects on fitness and health, for instance olfactory receptors. In contrast, rare predicted LOF genotypes in these outbred samples were usually heterozygous and thus of uncertain overall impact on gene function. Several approaches have been described recently to identify naturally occurring human knockouts. A large exome sequencing aggregation study (ExAC), of predominantly outbred individuals, identified 1,775 genes with homozygous predicted LOF genotypes in 60,706 individuals(*2*). In population isolates, 1,171 genes with complete predicted LOF were identified in 104,220 Icelandic individuals(*3*), and modest enrichment for homozygous predicted LOF genotypes shown in Finnish individuals(*4*). However even in these large samples homozygous predicted LOF genotypes tend to be associated with moderate (around 1%) rather than very rare allele frequencies, and these approaches will not readily assess knockouts in most genes even if they could be scaled up.

Here, we identify knockouts created by rare homozygous predicted loss of function (rhLOF) variants by exome sequencing 3,222 Pakistani-heritage adults living in the UK who were ascertained as healthy, type 2 diabetic, or pregnant **[supplementary materials section S1 (SM S1**)]. These individuals have a high rate of parental relatedness (often with parents who are first cousins) and thus a substantial fraction of their autosomal genome is homozygous. We extend our analysis to interpreting the clinical impact of the identified knockouts by linking to healthcare and epidemiological records, with the main aims of i) assessing the clinical actionability and health effects of naturally occurring knockouts in an adult population, ii) understanding the architecture of gene essentiality in the human genome, through the characterisation of the population genetics of LOF variants, and iii) studying in detail a particularly interesting knockout (*PRDM9*).

The average fraction of the coding genome in long homozygous regions inferred to be identical by descent from a recent common ancestor (“autozygous”) in our samples was 5.62%, much higher than that in outbred European heritage populations **(figs. 1 A, S4).** We identified per subject on average 140.3 non-reference sequence predicted LOF genotypes comprising 16.1 rare (minor allele frequency <1%) heterozygotes, 0.34 rare homozygotes, 83.2 common heterozygotes and 40.6 common homozygotes. Overall we identified 1,111 rhLOF genotypes at 847 variants (575 annotated as LOF in all GENCODE-basic transcripts) in 781 different protein-coding genes **(fig. 1, Data File S1, SM S2)** in 821 individuals. Nearly all rhLOF genotypes were found within autozygous segments (94.9%, **SM S3),** and the mean number of rhLOF per individual was proportional to autozygosity **(fig. 1B).** Autozygous segments were observed across all exomic sites with a density distribution not significantly different from random **(SM S3,** Kolmogorov-Smirnov P=0.112). Based on these values we estimate that 41.5% of individuals with 6.25% autozygosity (expected mean for individual with first-cousin related but otherwise outbred parents) will have one or more rhLOF genotypes.

**Fig. 1.**
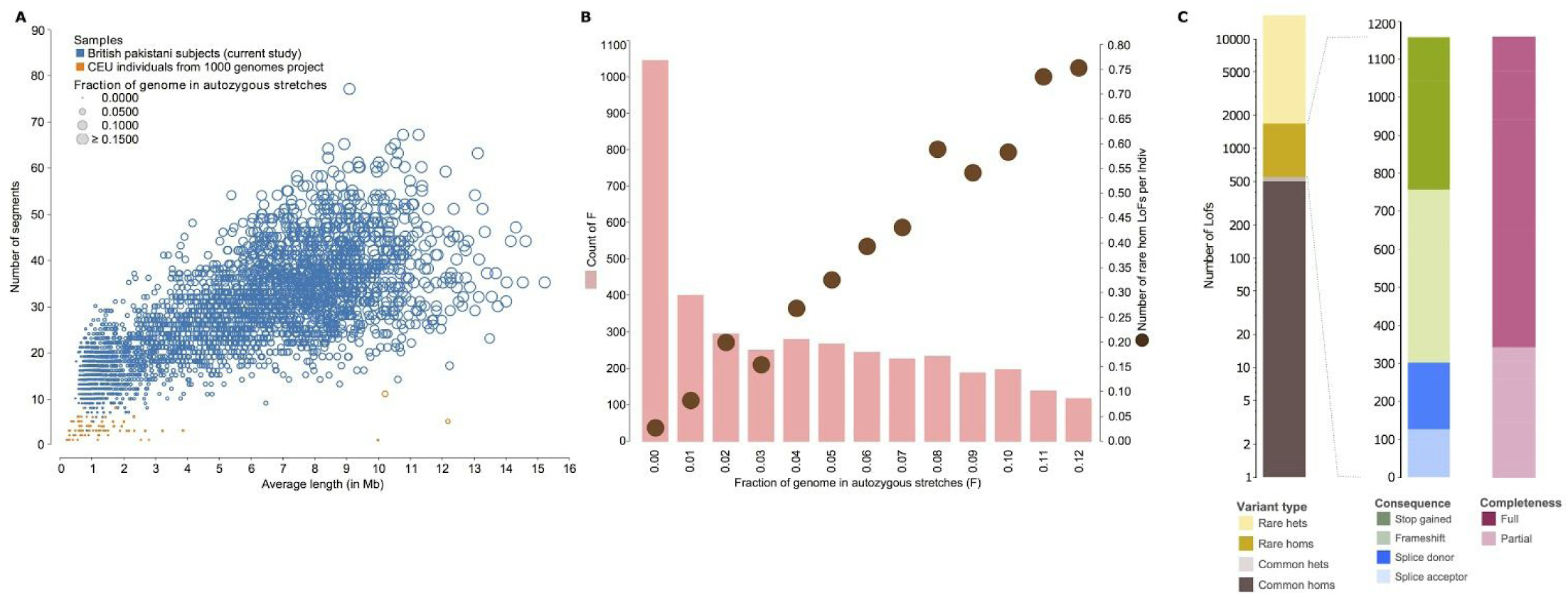
Discovery and annotation of rhLOF variants. **(A)** Autozygous segment numbers and length for Pakistani-heritage subjects in the UK, and 1000 Genomes project European (CEPH Utah residents with ancestry from northern and western Europe; CEU) individuals. **(B)** Autozygosity and rhLOF in 3,222 individuals. Count of number of individuals (left Y axis, pink columns) binned by fraction of autozygous genome (X axis, showing values from 0.00 to 0.12), with mean number of rhLOF genotypes per individual (right Y axis, brown circles). (C) Distribution of LOF variants by allele frequency, heterozygous or homozygous genotype, predicted protein consequence, and whether predicted for a full or partial set of GENCODE-Basic transcripts for the gene.

The majority of our rhLOF containing genes were novel, in that rhLOF genotypes for them had not been previously reported. We observed 167 rhLOF genes that were also reported as containing homozygous or compound heterozygous LOF genotypes in Iceland, and 299 rhLOF genes reported as containing homozygous LOF genotypes in ExAC. There were 107 rhLOF genes common to all three datasets **(SM S4, Data File S2)** suggesting a subset of genes either tolerant of LOF and/or with higher rates of mutation, and 422 rhLOF genes private to our dataset. We observed 89 rhLOF genotypes as homozygous LOF alleles without observing any heterozygotes, showing the ultra-rare genotypes identifiable in our study design but not in bottlenecked populations, and observed three subjects with 5 rhLOF genotypes (F=12.1 %, 16.7%, 23.8%). By downsampling the discovered heterozygous variation, we predict that in 100,000 subjects with related parents (F=6.25%) of the same genetic ancestry we would expect at least one knockout identified in 8,951 of the ~20,000 human protein-coding genes **(fig. S3, SM S5).** This estimate, taken together with the observed rhLOF discovery rate per person, demonstrates that the related parent design is a much more efficient strategy to identify large numbers of naturally occurring complete human knockouts, compared to sequencing all-comers or the population bottleneck design. Effectively all the knocked-out genes in the Icelandic population have now been discovered, whereas sequencing subjects with related parents will continue to yield new genes until all those in which human knockouts compatible with life are exhausted.

Next, we used the distribution of rhLOF and other variants seen in our sample to explore questions about recessive selection and gene function. We observed a significantly lower density of annotated rare LOF variants within autozygous tracts, where they are homozygous, compared to outside autozygous tracts, where they are typically heterozygous, whilst control synonymous variants showed no difference **(fig. 2A).** Since the genomic location of autozygous regions appears random, this likely reflects direct selection against rhLOF genotypes. We quantified this deficit by matching each of the 16,708 rare annotated LOF (heterozygous and homozygous) variants to a randomly selected synonymous variant of the same allele frequency, and observed that on average 975.5 of the matched rare synonymous variants were seen in a homozygous state compared to 842 rhLOF variants, indicating a deficit of 13.6% (95% confidence interval 8–20%) of rhLOF variants **(fig. 2B, SM S6).** The missing fraction is presumably not seen because they result in early lethality or severe disease incompatible with our selection criteria as healthier adults. This estimate of the fraction of gene knockouts essential for healthy life is lower than previous estimates of ~30% derived from knockout mouse models(*5*). However it is higher than the 6.4% deficit of below 2% frequency LOF alleles in the Icelandic population obtained from analysis of parent-child double transmissions(*3*), consistent with that analysis having excluded very rare alleles below 0.1% frequency and hence being biased towards mutations that had already been subject to selection.

**Fig. 2.**
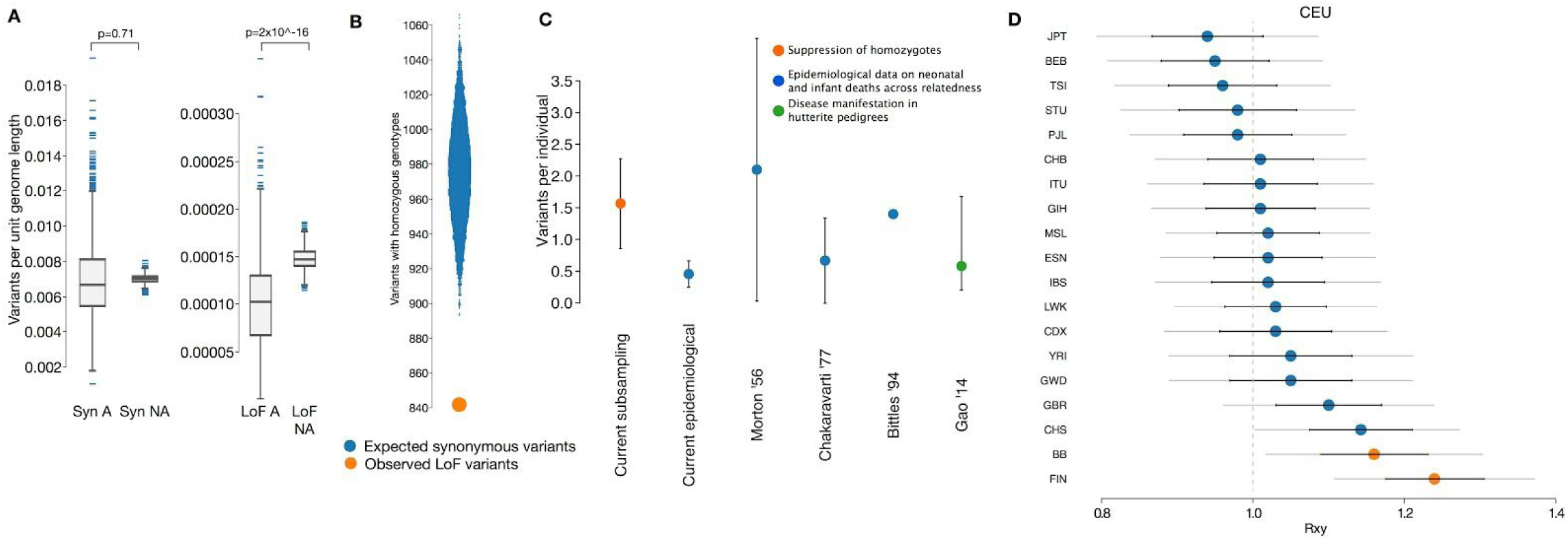
Population genetic analysis of rhLOF variants. **(A)** Comparison of number of LOF variants per unit length in autozygous regions (LOF A) with expected rate from non-autozygous sections (LOF NA) showing suppression of rhLOFs. A similar analysis of synonymous (Syn) variants shows no significant differences. **(B)** Observed number of variants with homozygote genotypes in 16,708 rare LOF variants (orange circle) versus a frequency matched subsampling of synonymous variants (blue violin plot). (C) Quantification of the number of recessive variants incompatible with healthy human adult life carried on average by a single individual. Direct subsampling estimate for rhLOF variants (orange circle) from current study; epidemiological estimates based on correlating infant mortality rates to estimated autozygosity (blue circles) in current and published data; direct estimate from large Hutterite pedigree (green circle). 95% confidence intervals as black bars. **(D)** Relative number of derived LOF alleles that are frequent in one population and not another (under neutrality the expectation is 1.0), calculated for 1000 Genomes Project populations and the current Birmingham/Bradford Pakistani heritage population (BB), compared to the CEU population. Error bars represent ±1 (black) or 2 (grey) standard errors, and significant differences versus CEU population are highlighted in orange circles.

We then combined the calculated deficit rate with the observed number of heterozygous annotated LOF variants, integrating across allele frequencies, to obtain a direct estimate of the recessive lethal load per person. This suggests that a typical individual from the population we sampled carries 1.6 recessive annotated LOF variants incompatible with adult life in the heterozygous state **(fig. 2C, SM S7).** Previous studies have estimated a related quantity by correlating the number of miscarriages, stillbirths and infant mortalities with the level of autozygosity(6, *7*). Using epidemiological data from **13,586** mothers from the same Born In Bradford birth cohort studied in our genetic analysis, we were able to perform a corresponding correlational analysis, and estimated **0.5** lethal equivalents per individual **(SM S7).** This is the largest such study to date, utilizes our direct genetic measurements of autozygosity, and is well controlled for socioeconomic factors. The difference between the two estimates can be accounted for by the fact that the first includes embryonic lethals, whereas the second only involves deaths after a registered pregnancy. This suggests there are twice as many recessive mutations that are embryonically lethal as those that result in foetal or infant death. The absolute values of the estimates for mutational load may be biased downwards in the population we are studying compared to outbred populations, due to purging of recessive lethal mutations by the long-term endogamous marriage structure. We examined the consequences of this structure by using a method previously applied to examine the consequences of population bottlenecks(*8*-*10*). Controlling for other effects by comparing to synonymous mutations, we see a significant but moderate increase in purging of the rhLOF mutational load in our Pakistani heritage population dataset compared to outbred populations from the **1000** Genomes Project, although less than that caused by the historic bottleneck in the Finnish population (FIN in **fig. 2D, SM S8).**

We examined 215 genes with rhLOF in our dataset that have an exact 1:1 mouse:human gene ortholog to mouse and mouse gene knockout data. Of these, there were 52 genes where a lethal mouse phenotype had been reported on at least one genetic background, confirming previous observations of differences in lethality between humans and laboratory mouse strains(*11*). We found that whether or not the mouse ortholog knockout is lethal is not associated with alteration of protein function, duplication or changes in gene expression **(SM S9).**

Previous work suggests that LOF variants might be enriched for sequencing errors(*1*). We found complete concordance between our exome sequencing genotype and Sanger dideoxy-sequencing genotype for 35 rhLOF genotypes (a further 2 could not be assayed) in samples from recalled individuals **(SM S10, table S1).** For a subset of genes known to be expressed in blood we observed the absence of protein on western blots using whole blood samples for 5 rhLOF genotypes (*LSP1, DPYD, GCA, SAMD9, MSRA*), weak protein (EMR2); and/or very low RNA expression compared to other samples for 6 rhLOF genotypes (*LSP1, SLC27A3, GCA, SAMD9, MSRA, EMR2*)(**SM S11, fig. S1).** Extensive validation of LOF variants has been described elsewhere using RNA sequencing(*3, 12*).

Our observed modest depletion of rhLOF in adult subjects, due to presumed deleterious effects, raises the question: what effect do the complete knockouts we observe have on clinical phenotypes? In subjects from the Born In Bradford study, where full health record data was available, we observed 55 rhLOF genotypes in 53 individuals in Online Mendelian Inheritance in Man (OMIM) confirmed recessive disease genes. We first carefully inspected the read alignments used to call genotypes, observing excellent genotype calls and no anomalies **(SM S12),** in all but one case, an indel in a short homopolymer tract with a rare single nucleotide variant 3bp 3’ (*GHRHR,* also excluded by annotation review). Next we inspected the annotation of these variants, suspecting enrichment for annotation errors(*1*). Taking a cautious approach, we considered 13 of 55 rhLOF genotypes to be possible genome annotation errors (i.e. incorrectly described as LOF): 2 genotypes where the GENCODE reading frame appeared to be mis-assigned (*CLN3*); 3 genotypes where the variant is most likely in the UTR due to mis-assignment of the reading frame of the last exon in GENCODE (*NF1, POMGNT1*); 2 genotypes for a splice-site indel where the Ensembl Variant Effect Predictor has likely mis-annotated as LOF (*C12orf57*); and 6 genotypes where the protein-coding status of the LOF transcript appears questionable (*LRTOMT, DPM2, GHRHR, DTNBP1, PUS1*) **(SM S12, table S2).** We note that these are all putative genome annotation, rather than sequencing, errors and this highlights the great care that must be taken in interpreting LOF genotypes predicted through exome or genome sequencing.

Surprisingly, for only six of these 42 (excluding all putative annotation errors) rhLOF genotypes were lifetime primary health record diagnoses **(SM S13)** definitely compatible with OMIM phenotype (*FLG, NPC2, TRPM1, DNAI2, DYSF*), with a further three genotypes suggestively compatible (*GJB4, NME8, ARMC4*)(**table S3).** This suggests a greater extent of incomplete penetrance; or unrecognised technical issues with sequencing or annotation (see above); or considerable incorrect OMIM gene-phenotype assignment (i.e. incorrect assignment in the original published report(s)). Our study has an ascertainment bias in that subjects were recruited as adults, either as pregnant women attending antenatal clinics, healthy participants from the general population, or for a type 2 diabetes study. This is the opposite end of the health spectrum to patients with severe and young-onset diseases (often from multiply affected families) who comprise much of the Mendelian disease literature. We recognize that health record analysis identifies symptoms and conditions that present to medical services, and as such differs from bespoke phenotyping with prior knowledge of genotype (e.g. audiometry for deafness genotypes). However in support of the general significance of our findings, markedly reduced penetrance estimates have been shown for Mendelian forms of diabetes when taken from population samples(*13*), and the ClinGen study identified an initial set of variants annotated as pathogenic in OMIM (220 of~25,000 entries) but now thought to be benign or of uncertain significance(*14*). In the ClinSeq cohort, with an older age of ascertainment, 45 of 73 individuals with rare heterozygous LOF in a gene that had been reported to cause disease via dominant LOF alleles did not have the expected phenotype(*15*). The UK10K study also suggested that reported estimates of disease-variant penetrance were too high(*16*). These findings have increasing relevance as exome and genome sequencing move into population-scale surveys - for example we observed adults with a *GJB2* knockout (p.Trp77Ter, a published deafness risk variant(*17*)) without any symptoms of hearing loss, and a *MYO3A* knockout without any symptoms of hearing loss and with normal audiometry(*18*, *19*).

We next assessed primary care electronic health records to look for more general effects of rhLOF genotypes on health status in Born In Bradford adults since study recruitment **(SM S14).** Drug prescription rate and clinical staff consultation rate have been shown to correlate strongly with health status(*20*). We compared individuals with one or more rhLOF (n=638) to individuals without rhLOF (n=1524), and found no association with prescription rate (OR 1.001, 95% Cl 0.988 – 1.0144) or consultation rate (OR 1.017, 95% Cl 0.996 – 1.038). Nor did we find association with prescription rate (OR 1.006, 95% Cl 0.966 – 1.045) or consultation rate (OR 1.029, 95% Cl 0.964 – 1.090) in individuals (n=53) with rhLOF in OMIM confirmed recessive disease genes compared to individuals without rhLOF; nor on prescription rate (OR 1.005, 95% Cl 0.968 – 1.043) or consultation rate (OR 1.038, 95% Cl 0.978 – 1.094) in individuals (n=58) with rhLOF in mouse knockout lethal genes. Findings were unchanged in additional adjusted analyses including age, education, duration of available data, and autozygosity.

To illustrate the potential of our human knockout discovery approach, we identified a healthy adult woman with a *PRDM9* rhLOF genotype (p.R345Ter in the sole GENCODE-basic transcript, arising within a 25Mb autozygous region), and we confirmed her genotype by Sanger sequencing **(SM S15, fig. S6A, S6B).** No human knockout of *PRDM9* has been described previously. DNA was available from one of her three healthy children who was confirmed heterozygous, as expected for germline transmission, for the same variant **(fig. S6B).** Somatic but not germline homozygosity resulting from a crossover early in embryonic development leading to somatic loss of heterozygosity (LOH) is excluded by the fact that this subject is heterozygous on both sides of the autozygous region. Should LOH have occurred, she would be entirely homozygous from the site of the originating crossover to the telomere **(fig. S6C).** Review of her lifetime primary and secondary care health records was unremarkable. Her genotype predicts protein truncation in the SET methyltransferase domain (thus lacking the DNA-binding zinc-finger domain), which was confirmed in an *in vitro* expression system(**SM S15, fig. 3).**

**Fig. 3.**
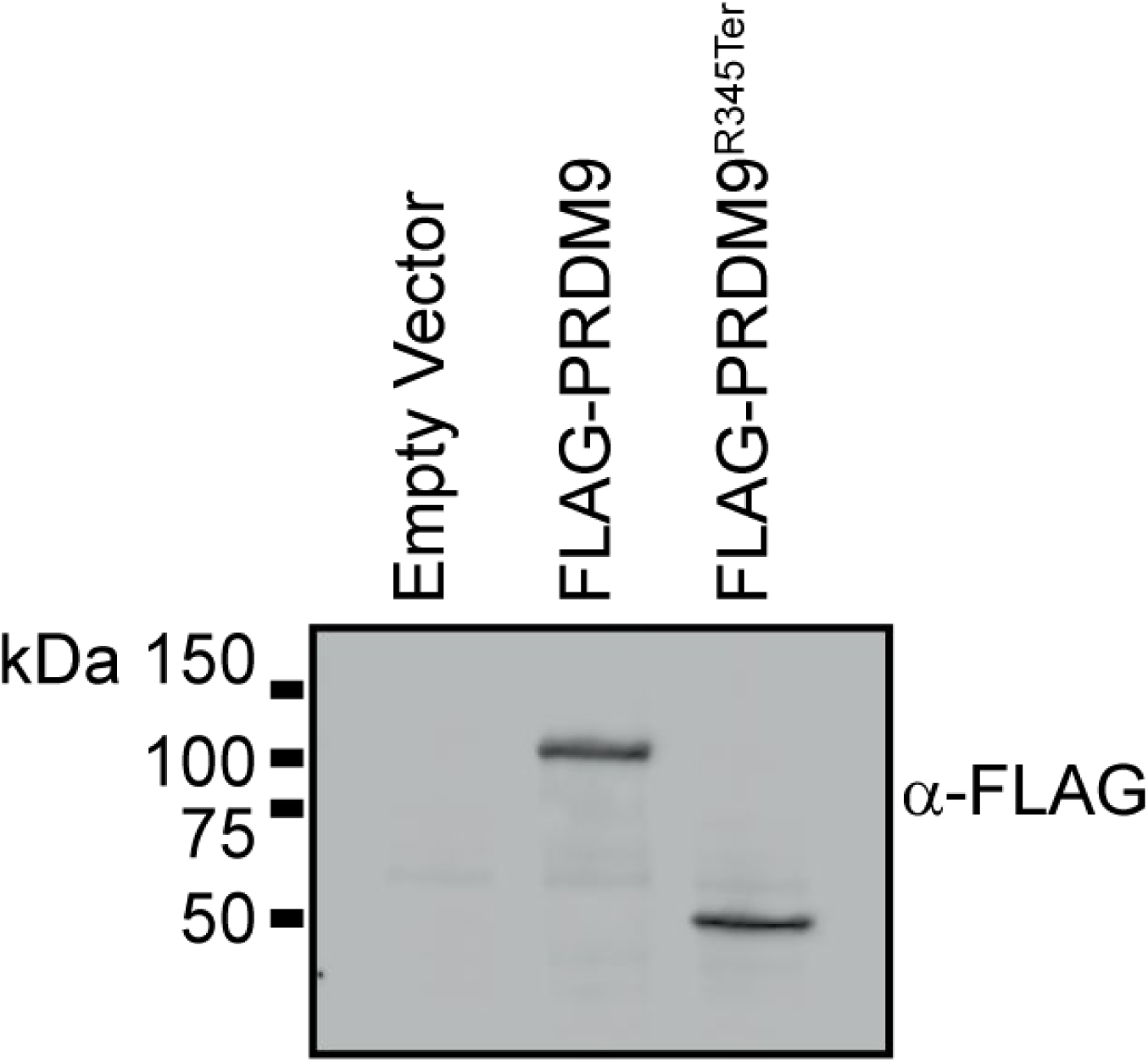
Expression of PRDM9 in HEK293 cells. FLAG-tagged *PRDM9* constructs were expressed for 24h in HEK293 cells. The expected protein sizes (full length 110.1 kDa, R345Ter 43.2kDa) were confirmed on western blot.

*PRDM9* is the major known determinant of the genomic locations of meiotic recombination events in humans and mice(*21–23*). *PRDM9* is highly polymorphic in humans, and the common European *PRDM9-A* allele (for which the knockout subject is homozygous) and the common African *PRDM9-C* alleles activate very different sets of meiotic recombination hotspots(*24, 25*). To determine the sites of meiotic recombination in the maternal gamete transmitted from the PRDM9 R345Ter knockout mother to her child we used a novel approach of long-range directly-phased whole-genome sequencing developed by 10XGenomics, identifying 39 candidate crossovers **(SM S15).** As a control, we sequenced a widely-studied mother-child duo homozygous for the *PRDM9-A* allele (NA12878 / NA12882 CEPH individuals from the Coriell repository) using the same technologies **(DataFile S3).** High specificity of this approach was shown by excellent concordance (40/42) between our observation of 42 recombination sites, versus a set of 54 recombination sites generated by standard methods using published genotypes from 11 children, parents and grandparents from this large CEPH pedigree.

Most meiotic double-strand DNA breaks (DSBs) occur at discrete sites determined by the DNA binding specificity of PRDM9 which localises histone 3 lysine 4 tri-methylation (H3K4me3), although genetic crossover is only one outcome of DSB repair. Human linkage disequilibrium maps are a superimposition of *PRDM9* allele specific DSB maps(*25*). Using DSB maps and a maximum likelihood model **(SM S15)** to account for variability in region size and hotspot density(*22*), we estimated that only 5.9% (2 log unit confidence interval: 0 – 24%) of the observed *PRDM9* knockout duo maternal gamete crossovers matched DSB sites. In contrast, 52.1% (confidence interval: 36 – 69%) of the NA12878 / NA12882 duo crossovers occurred in known *PRDM9-A* allele DSB hotspots. As a second line of evidence we show that 18.5% of crossovers observed in the *PRDM9* knockout duo (confidence intervals: 1% – 42%) occurred in linkage disequilibrium based hotspots versus 75.7% (confidence interval 57%-89%) in the NA12878 duo **(SM S15).** Our figures for the NA12878 / NA12882 duo are consistent with previously published estimates of 60% (confidence interval: 58%-61%) from multiple individuals(*22*). *Prdm9* knockout mice demonstrate abnormal location of recombination hotspots with enrichment at gene promoters and enhancers, and also fail to properly repair double-stranded breaks and are infertile (both sexes sterile)(*26*, *27*). Dogs, the only known mammalian lineage lacking *Prdm9,* lost early in canid evolution, retain recombination hotspots which unlike humans or knockout mice occur in high GC content regions(*28*). It has been speculated that dog recombination is controlled by an ancestral mammalian mechanism, and that PRDM9 competes and usurps these sites when active in non-canids(*28*, *29*). However we did not see an increased overlap in our PRDM9 R345Ter duo crossover intervals with promoters and their flanking regions from ENSEMBL, or with H3K4me3 maps from multiple cell types, or with high GC content, compared to the NA12878 duo control **(SM S15).** Our identification of a healthy and fertile *PRDM9*-deficient adult human suggests differences from mouse, and supports the possibility of alternative mechanisms of localising human meiotic crossovers(*25*).

In drug discovery, genetic information improves the success of phenotype-target validation(*30*), for example the development of PCSK9 inhibitors relied on causal genotype-phenotype effects including both gain- and loss-of-function. We observed no difference in the proportion of genes with known or predicted druggable targets for rhLOF genes (15%) compared to all genes (13%, P=0.098) **(SM S16).** However the rhLOF gene drugged targets have a higher than expected development success, with 11.4% phase I to approval rate versus 6.7% for all target-indication pairs (P=0.046). The observation of rhLOF in a drug target gene in a healthy individual provides an indication that the gene (or its protein product) might be safely knocked down.

We investigated the protein-protein interaction properties of observed rhLOF genes, and compared this with the Icelandic LOF dataset, and with a catalogue of rare disease genotype-phenotype associations (Orphanet) further divided into loss- or gain-of-function genotypes **(SM S17).** Genes containing rhLOF showed 50% fewer molecular interactions compared to all genes in the STRING interactome dataset (P=3.4 ×10^−9^), predominantly driven by the Binding Interaction class (P=9.3 ×10^−11^). Similar effects were seen in the Icelandic data **(table S4),** and this was in contrast to both known pathogenic LOF variants and especially pathogenic gain-of-function (GOF) variants reported in Orphanet, which showed the opposite effect of increased overall molecular interactions (P=1.1×10^−6^, 2×1 O'^12^ respectively), driven by increased interactions across all interaction classes. The reduced protein-protein interactivity of rhLOF genotypes observed in our generally healthier cohorts may contribute to the observed drug clinical development success, and to the relatively few effects on clinical phenotypes and general health measures.

Taking together these data, we suggest that apparent rhLOF genotypes identified by exome or genome sequencing from adult populations will require cautious interpretation. This variant class has the greatest predicted effect on protein function, nonetheless loss of some proteins is harmless to the individual, and LOF variants may not always be as clinically actionable as previously considered. The challenge is to determine the balance between genome annotation errors, incorrect Mendelian disease gene-phenotype assignments, and true incomplete penetrance (the variability of mutational phenotypic consequences in different genetic backgrounds) - which will require very large studies. This becomes of increasing importance now that exome and genome sequencing is rapidly expanding into healthier adults and other non-disease individuals. Much remains unknown regarding clinical interpretation of, and attribution of causality for, LOF variants.

Finally, we suggest that efforts to identify naturally occurring human knockouts - whether in bottlenecked populations, or more efficiently in subjects with related parents - will yield new biological insights, as exemplified by our identification here of a *PRDM9* deficient healthy and fertile woman.

## Acknowledgments

The study was funded by the Wellcome Trust (WT102627 & WT098051), Barts Charity (845/1796), Medical Research Council (MR/M009017/1). This paper presents independent research funded by the National Institute for Health Research (NIHR) under its Collaboration for Applied Health Research and Care (CLAHRC) for Yorkshire and Humber. Core support for Born in Bradford is also provided by the Wellcome Trust (WT101597). VN was supported by the Wellcome Trust PhD Studentship (WT099769). DGM and KK were supported by the National Institute of General Medical Sciences of the National Institutes of Health under award number R01GM104371. ERM is funded by NIHR Cambridge Biomedical Research Centre. Born in Bradford is only possible because of the enthusiasm and commitment of the Children and Parents in BiB. We are grateful to all the participants, health professionals and researchers who have made Born in Bradford happen. We thank Ms Beverley McLaughlin (QMUL) for assistance, and Dr Jeremy Rogers (HSCIC) for advice. We would like to thank the Exome Aggregation Consortium and the groups that provided exome variant data for comparison. A full list of contributing groups can be found at http://exac.broadinstitute.org/about.

RD declares his interests as a founder and non-executive director of Congenica Ltd., that he owns stock in illumina from previous consulting and is a scientific advisory board member of Dovetail Inc. MSL and KG are employees of 10XGenomics Inc.

Data reported in the paper are presented in the Supplementary Materials, and at the European Genotype-phenome Archive (www.ebi.ac.uk/eaa’) under accession numbers EGAS00001000462, EGAS00001000511, EGAS00001000567, EGAS00001000717 and EGAS00001001301.

### Supplementary Materials

Materials and Methods (SM S1 to S17)

Figs. S1 to S6

Tables S1 to S8

References (31–63)

### Other Supplementary Material for this manuscript includes the following

Data File S1 as an Excel file

Data File S2 as an Excel file

Data File S3 as an Excel file

### Materials and Methods

#### SM S1 Exome sequencing of 3222 Pakistani-heritage adult individuals living in the UK

##### S1.1 Subjects and phenotypes

Birmingham adult Pakistani-heritage subjects comprised 130 subjects from a Birth Cohort study in Birmingham, UK(*31*), 471 adult healthy, and 892 adult type 2 diabetes subjects from Birmingham and Coventry as part of the UK Asian Diabetes Study(*32*) - a total of 1493 DNA subject samples, resulting in **(after all quality control steps) 1060 exome sequenced subject samples.** Approval was from the Birmingham East, North and Solihull Research Ethics Committee and from the South Birmingham Research Ethics Committee.

Born in Bradford (BiB) is a longitudinal multi-ethnic birth cohort study aiming to examine the impact of environmental, psychological and genetic factors on maternal and child health and wellbeing(*33, 34*). The full BiB cohort recruited 12,453 women during 13,776 pregnancies between 2007 and 2010. Ethical approval was granted by Bradford Research Ethics Committee. BiB adults studied here comprised 2490 DNA subject samples from pregnant women (>95% self-stated Pakistani-heritage) ascertained at an antenatal clinic visit in Bradford, UK(*33*), resulting in **(after all quality control steps) 2162 exome sequenced subject samples.** The first 1570 samples were of unselected parental relatedness status and included 554 with self-stated first cousin parental relatedness, the remaining samples were all of self-stated first cousin parental relatedness. We deliberately included (before the quality control stage) duplicate samples (taken at different antenatal visits for each of multiple pregnancies). We also recalled **(SM S10)** selected subjects for a second sample for genotype validation by a different method.

##### S1.2 Exome sequencing and sequence-level quality control

Genomic DNA was extracted from peripheral blood, and quality confirmed by agarose gel electrophoresis (requiring high molecular weight) and picogreen assay. Samples were genotyped for identity checking by either Sequenom assay (26 common autosomal variants, and 4 gender markers) or Fluidigm assay (22 common autosomal variants, and 4 gender markers). Samples where DNA-based gender and stated gender were mis-matched were excluded.

Genomic DNA (approximately 1 ug) was fragmented to an average size of 150 bp and subjected to DNA library creation using established Illumina paired-end protocols. Adapter-ligated libraries were amplified and indexed via PCR. A portion of each library was used to create an equimolar pool comprising 8 indexed libraries. Each pool was hybridised to SureSelect RNA baits (Human All Exon V5, Agilent Technologies) and sequence targets were captured and amplified in accordance with manufacturer’s recommendations. Enriched libraries were used for 75 base paired-end sequencing (HiSeq 2000, Illumina) following manufacturer’s instructions. Sequencing was performed to an expected ~40× read-depth, since the primary aim was to identify homozygous genotypes in autozygous regions.

##### S1.3 Sample filtering

Stated duplicates were obtained from the BiB cohort, which took repeat samples when the same individuals registered at the Bradford hospital maternity ward for multiple pregnancies. A total of 149 individuals who registered for 2 pregnancies and 4 individuals who had registered for 3 pregnancies were recorded. From these duplicate pairings a total of 153 pairs had sequence data. Duplicate checks were performed using bcftools (version: 1.1–113-gd991f3f used throughout) gtcheck-G1 to compute the pairwise discordance between samples. 1000 random bi-allelic SNP sites were chosen out of a set of markers that were of at least MAF>0.05 in both the BiB dataset as well as the 1000 Genomes Project Phase3 release set, and had at least a mean sequencing depth of 20. Duplicates were clearly separated from other samples in a plot of number of discordant genotypes between each pair of sample (not shown). A duplicate sample was then removed from each pair so that there were unique samples for subsequent analyses.

To confirm sample identity and exclude laboratory mix-ups we compared the genotype calls made using Sequenom or Fluidigm as described above with those from our sequencing data. We removed 30 samples for which there was a genotype discordance of more than 30%.

Principal components analysis (PCA) to determine individual ethnicity was performed by merging bi-allelic SNPs from the current dataset with the 1000 Genomes Project phase 3 release set. Bi-allelic SNP sites that were of at least MAF>0.05 in each of the datasets were taken to define a set of markers used to perform the PCA analysis. PCA was first performed with 1000 Genomes Project data using all populations, and samples from the current dataset projected onto that reference. Pakistani-heritage subjects could clearly be defined, distinct from other South Asian ancestries on component 2 and 3 of the PCA and a region of the plot containing the majority of the samples was defined, leading to the removal of 294 samples that plotted outside the region **(fig. S2).** These 294 contained 20 samples with a 10-fold enrichment in the number of singleton sites which support ancestry other than the main cohort of British Pakistani subjects. Finally, using VerifyBamID(*35*) (freemix parameter > 0.03), we removed 24 samples that were predicted to have contamination between the samples.

##### S1.4 Alignment and BAM processing

Following generation of raw reads, these were converted to BAM format using illumina2BAM and lanes de-multiplexed so that the tags were isolated from the body of the read, decoded, and used to separate out each lane into lanelets containing individual samples from the multiplex library and the PhiX control. Reads corresponding to the PhiX control were mapped and used with WTSI's spatial filter program to identify reads from other lanelets that contained spatially oriented INDEL artefacts and mark them as QC fail. Reads were aligned with BWA-MEM to the GRCh37+decoy reference genome used by the 1000 Genomes project (ftp://ftp.1000genomes.ebi.ac.uk/vol1/ftp/technical/reference/phase2referenceassemblyseauence/hs37d5.fa.az’). PCR and optically duplicated reads were marked using Picard MarkDuplicates, and after manual QC passing data was deposited with the EBI-EGA.

In order to ensure the quality of the large quantity of BAMs produced for the project, an automated quality control system was employed to reduce the number of data files that required manual intervention. This system was derived from the one originally designed for the UK1 OK project(*16*) and used a series of empirically derived thresholds to assess summary metrics calculated from the input BAMs. These thresholds included: percentage of reads mapped; percentage of duplicate reads marked; various statistics measuring indel distribution against read cycle and an insert size overlap percentage. Any lane that fell below the "fail" threshold for any of the metrics were excluded; any lane that fell below the "warn" threshold on a metric would be manually examined; and any lane that did not fall below either of these thresholds for any of the metrics was given a status of "pass" and allowed to proceed into the later stages of the pipeline.

Passed lanelets were then merged into BAMs corresponding to sample's libraries and duplicates were marked again with Picard after which they were then merged into BAMs for each sample. Next, for each sample we re-aligned reads around known and discovered indels followed by base quality score recalibration using GATK (version: v3.3–0 used throughout). Lastly samtools ‘calmd’ was applied and indexes were created. Known indels for realignment were taken from Mills-Devine(*36*) and 1000 Genomes Project Phase 1 (*37*) low coverage set, available from the 1000 Genomes ftp site. Known variants for base quality score recalibration were taken from dbSNP 137.

##### S1.5 Variant calling

Two variant call-sets, one with the Genome Analysis Toolkit(*38*, *39*) (GATK) HaplotypeCaller and one with samtools/bcftools(*40*), were made from the 4,353 samples that passed QC measures to this point. Calling was restricted to the Agilent V5 exome bait regions +/- a 10Obp window on either end. The following parameters were used:

*GATK HaplotypeCaller* [GATK version: v3.3–0]:

Single-sample genome VCF (gVCF) files were created using GATK HaplotypeCaller run in gVCF mode on each of the sample BAM files using parameters:

-T HaplotypeCaller -R hs37d5.fa -variant_index_type LINEAR -variant_index_parameter 128000
--disable_autoJndex_creation21478889_and_locking_when_reading_rods -minPruning 3
--maxNumHaplotypeslnPopulation 200 -ERC GVCF -max_alternate_alleles 3 -contamination 0.0
-L Agilent_human_exome_v5_S04380110/S04380110_Covered.baits.nochr.wl 00.nr.bed

For each chromosome, we ran CombineGVCFs in batches of~65 samples using parameters:

-T CombineGVCFs -R hs37d5.fa -disable_autoJndex_creation_and_locking_when_reading_rods
--variant samplel.vcf.gz -variant sample2.vcf.gz … -isr INTERSECTION
-L $chr -L Agilent_human_exome_v5_S04380110/S04380110_Covered.baits.nochr.wl 00.nr.bed

Then, for each chromosome, we ran GenotypeGVCFs on the output of CombineGVCFs using parameters:

-T GenotypeGVCFs -R hs37d5.fa -disable_autoJndex_creation_and_locking_when_reading_rods
--variant groupl.vcf.gz -variant group2.vcf.gz … -isr INTERSECTION
-L $chr -L Agilent_human_exome_v5_S04380110/S04380110_Covered.baits.nochr.wl 00.nr.bed

*SAMtools/BCFtools*: [samtools version: 1,1–30-g7f47a7c, bcftools version: 1.1–113-gd991 f3f, htslib version: 1,1–104-g948a68c]:

All-site, all-sample BCF files from all the BAM files were generated using 'samtools mpileup'. This processing was split into 100Mb chunks across the genome using parameters:

samtools mpileup -t DP,DPR,INFO/DPR -C50 -pm3 -F0.2 -d2000 -L500 \
-I Agilent_human_exome_v5_S04380110/S04380110_Covered.baits.nochr.wl 00.nr.bed \
-g -r $chr:$from-$to -b $bam_list -f hs37d5.fa > $chr:$from-$to.bcf

For each chunk, variants were called using `bcftools call´ using parameters:

bcftools call -vm -f GQ $chr:$from-$to.bcf | bgzip -c > $chr:$from-$to.vcf.gz

On chromosome X, male samples were treated as diploid in the pseudo-autosomal regions (X:60001–2699520 and X:154931044–155270560) and haploid otherwise using the '-X' option in 'bcftools call'.

From the two initial callsets produced using GATK and samtools, we created a consensus call-set by intersecting the set of bi-allelic sites produced by each caller. The concordance between the two call-sets for SNPs was 95%, discordant genotypes were set to missing, variants with >1 % missing genotypes were excluded, and all subsequent analyses performed on this consensus call-set. The ratio of transitions to transversions (Ts/Tv) in the consensus call-set was 2.52 for all variants and 2.42 for singletons (37.3% of SNPs), with much lower Ts/Tv ratios for discordant calls.

We did not attempt to identify large deletions (e.g. >100bp) in our dataset, as these methods remain inaccurate for medium depth sequencing (the ~40x depth used here for most samples); we note that as a consequence we will have called a hemizygous LOF genotype (with the other allele a deletion) as a homozygote, but that this would still lead to complete allelic inactivation.

##### S1.6 Estimating variant quality

Estimation of error rates in exome sequencing data typically includes comparison with different technologies, or pedigree analysis, which whilst effective may miss systematic errors. A recent study(*41*) used a haploid human cell line, reporting heterozygous calls in the data as a proxy for error rate. Here, we use a similar strategy by examining the number of heterozygous calls within long autozygous stretches in our final call-set of 3222 Pakistani-heritage adults. We expect that long stretches of >10Mb autozygosity have occurred due to a recent inbreeding event, and therefore we can attribute heterozygous calls within such regions to either a de-novo mutation/gene conversion event, or a sequencing error. As the regions of autozygosity across these individuals occur due to random recombination events, we are able to measure errors occurring on multiple haplotypes and on joint called data.

We restricted autozygous regions considered to those at least 10Mb long and with no other region in that individual starting within 3Mb, and excluding the terminal 1Mb of each autozygous region. These stretches are long (often comprising a fifth of an entire chromosome) and heterozygous calls within them are equally likely to be seen in the middle of such stretches as compared to the ends (t.test p-value = 0.74). Overall we saw 88192 heterozygote calls in 12676437517bp of autozygous callable exome sequence, a rate of less than 1 per 100,000bp. Downsampled to the length of a single genome, this amounts to 19831 false heterozygotes per genome, within the range previously reported of 15000–30000 false heterozygotes per haploid genome sequenced at high coverage without PCR artifacts(*42*). This is still much higher than the expected rate due to new mutations, which is approximately 10^−7^ per bp (6 generations at a mutation rate of 1.5*10^−8^ per bp per generation), so most heterozygote calls within the autozygous sections are due to errors.

##### S1.7 Final variant filtering

We next used the Variant Quality Score Recalibration (VQSR) tool within GATK to calibrate the probability of variant call error, and further filter the dataset based on this single estimate for the accuracy of each call. We trained the VariantRecalibrator Gaussian mixture model using a set of true sites from HapMap project variants(*39*). We used the following metrics to train the model - for SNPS: QD, FS, MQRankSum, ReadPosRankSum, BaseQRankSum, MQ, InbreedingCoeff, SOR, GQ_MEAN, NCC; and for indels QD, FS, ReadPosRankSum, BaseQRankSum, MQ, InbreedingCoeff, SOR, GQ_MEAN, NCC. After calibrating using the heterozygotes in autozygous regions identified above as false positives **(fig. S5),** we chose the VQSR 99% filtering threshold for SNPs, and no VQSR filtering on the indels (VQSR did not increase specificity for indels). We used this as the **final dataset reported in the main results section of this paper.** In this final dataset, the SNP heterozygous call error rate is 1.47% (the rate of false heterozygote calls in autozygous regions as a fraction of the total rate of heterozyote calls in non-autozygous regions) and the corresponding indel heterozygous call error rate was 1.63%. The ratio of transitions to transversions (Ts/Tv) in the final call-set was 2.56 for all variants and 2.50 for singletons.

As an additional method to estimate error rate, we used 176 pairs of known duplicate samples (independent blood samples taken from Born In Bradford mothers at separate pregnancies). By examining the replication rate of heterozygous calls within autozygous sections in these individuals, we can classify our calls into those that are concordant; those likely to be due to systematic reasons (such as read mis-alignment to the genome, or de novo mutations) and those that are discordant; which are likely to be due to random issues in the sequencing process or sampling of reads during variant calling. For SNPs, there were on average approximately 27500 homozygous alternate calls/individual of which 1650 were in autozygous regions, with 99.25% replication in duplicate samples (99.6% in autozygous regions). With VQSR at 99% we lost 1.3% of these 27500 calls. There was a mean of 20 false heterozygote SNP calls/individual, with 50.1% replication in duplicate samples. For indels, there were mean 2068 homozygous alternate calls per person of which mean 123 were in autozygous regions, with 91.8% replication in duplicate samples (92.9% in autozygous regions). There was a mean of 2.6 false heterozygote indel calls/individual, with 24.8% replication in duplicate samples **(Table S5).** Using the 176 pairs of duplicate DNA samples from individuals to estimate the reproducibility of homozygous alternate allele genotypes in the final call set, we found a 0.5% SNP homozygous genotype discordance rate (0.3% within autozygous segments) and a 8.2% indel homozyous genotype discordance rate (7.1% in autozygous regions).

##### SM S2 Functional annotation of variants from exome sequencing

Loss-of-function (LOF) annotation was performed using the Loss-Of-Function Transcript Effect Estimator (LOFTEE, version 0.2, available at https://github.com/konradik/loftee’) a plugin to the Ensembl Variant Effect Predictor (VEP, version 77) based on GENCODE version 19 for the GENCODE basic set(*43*). LOFTEE considers all stop-gained, splice-disrupting, and frameshift variants, and filters out many known false-positive modes, such as variants near the end of transcripts and in non-canonical splice sites, as described in the code documentation. As a variant may have multiple different effects on different transcripts, the annotation of function is based on the most severe consequence per variant in the order as defined in Ensembl (http://www.ensembl.org/info/genome/variation/predicteddata.html.

##### SM S3 Identification of autozygous genomic segments

We applied a hidden Markov model (HMM) first utilized in(*44*) (https://samtools.github.io/bcftools/bcftools.html’) to identify regions of homozygosity (absence of heterozygous variation) due to parental relatedness with the important addition of utilising the fine scale sex averaged human recombination map(*45*). The allele frequency information was obtained using all 3,222 exomes and the transition parameters between autozygous and non-autozygous sections were learnt from the data using a Viterbi training scheme (segmental k-means algorithm), with the initial probabilities of being in the N (non-homozygous) or H (homozygous) state at the start of each chromosome set to equal. The resulting state assignments given by the Viterbi sequence with the optimal parameters comprised our inferred homozygous and non-homozygous tracts. All regions of homozygosity identified were >10kb in length.

The command line used for the inference using bcftools roh was: bcftools roh -G30 -a1e-8 -H1e-8 -e - -G30 -V -m genetic_map_chr{CHROM}_combined_b37.txt

We then compared our estimates of genetic autozygosity with those from self-stated estimates **(fig. S4A).** This confirmed our results in two ways. First, the estimate of genetic autozygosity corresponded with those from theoretical predictions from pedigree data, with offspring with higher parental relatedness having higher autozygosity. Second, we found that the median autozygosity from our genomic estimates was elevated over what we would expect in otherwise outbred individuals (first cousins, 6.25% and second cousins, 3.125%). This can be attributed to longstanding endogamy in the population which would lead to additional historic identity by descent. Additionally we noted that the variance even within individuals who stated that they were children of first cousins is extremely large, with estimates ranging from 0 to 25%.

After removing artifacts near locations close to the centromeres where we had low coverage in the sequencing data, we calculated the number of individuals that are autozygous at every site across the genome **(fig. S4B),** from which we can draw the following conclusions.

1. Every position in the genome contains at least one individual who is autozygous at a certain site, with a mean of 210.
2. The distribution of individuals who are autozygous at a site is not significantly different from random (Shapiro-Wilks test, see main text). The expectation was calculated by taking a weighted average of the overall inbreeding coefficient of each individual and assuming that the distribution of such sections would follow a binomial distribution and by approximation normal across 3,222 samples. Furthermore, the fact that these segments are randomly located across the genome and that such segments lie in hundreds of individuals with differing haplotypes means heterozygote sites can be used as a means of quality control of variants without any bias related to genomic location (SM1.6).
3. We find that 94.9% of all rhLOFs lie within the identified autozygous sections. This confirms that the overwhelming majority of homozygous genotypes were inherited from the parents at conception and hence not mosaics, ruling out mosaicism as a possible major reason for incomplete penetrance.

##### SM S4 Exome Aggregation Consortium (ExAC) and Icelandic population comparisons

We compared genes containing rhLOF in our dataset versus the genes containing homozygous LOF (restricting to high confidence variants including PASS all filters, and 80% call rate across all samples) in the Exome Aggregation Consortium (ExAC, http://exac.broadinstitute.org. data release 0.3 with VEPv79 annotation, accessed June 2015). A total of 1775 of 26915 genes in ExAC from 60706 individuals contained homozygous LOF genotypes. We included variants of all allele frequencies, as ExAC comprises multiple diverse ethnicities.

We compared genes containing rhLOF in our dataset versus the 1,171 genes containing homozygous LOF or compound heterozygous LOF for variants with a minor allele frequency <2% in the Icelandic sequenced dataset, using both direct sequenced and imputed Icelandic population data from 104,220 samples.

##### SM S5 Expected number of knockout variants to be seen in larger cohorts

The sequence data obtained from 3,222 healthy individuals provides us with a sample of the number and diversity of LOF variants (in both heterozygous and homozygous states) available in this specific population and we can use this to obtain estimates on the number of knockouts we expect to see in future sequencing up to 100,000 samples. In 3,222 individuals we observe 6,444 copies of each gene sampled from the population. In 100,000 people of first cousin offspring (each individual having 6.25% identity by descent), we expect to see 6250 homozygosed copies of each gene. Since this number is smaller than the number of observed copies (6,444) we could downsample the observed data set of haplotypes. As this does not account for heterozygous variation that would be incompatible with healthy life when homozygozed, we then reduce the estimated number of variant sites seen by 13.6% as this is the expected depletion of LOFs estimated using our subsampling approach **(SM S9).** We plot the results as a function of sample size in **fig. S3.** We expect these to be conservative estimates because we are including the regions that are autozygous in these individuals in our initial sampling. Based on this current data we note that there are a large number of genes yet undiscovered in which knockouts are likely to be compatible with adult human life, and that the discovery rate of these genes does not appear to plateau even if a study were to sequence 100,000 subjects with closely related parents. We also expect different sets of variation in different ethnicities and populations.

##### SM S6 Evidence for the effect of demography on natural selection in humans within a single generation

###### S6.1 Selection signatures of LOF variants in autozygous and non-autozygous regions.

We examined direct selection on recessive deleterious LOF variants by comparing the rates at which we observe variant sites of different mutational classes within and outside of autozygous tracts within individuals. Since the individuals ascertained in this study were healthier adults when sampled, recessive LOF variants that lead to death as an embryo, foetus or child, or result in severe early-onset Mendelian disease should be less missing from autozygous tracts, where they are exposed, while still present in non-autozygous portions of the genome. To control for the differing amount of autozygosity within the individuals as well as any differences in the relative rates of diversity inside and outside of autozygous sections, we normalize the total number of LOF genotypes seen in each section by the total number of variants in each section. We assess significance of the difference between relative rates of variants in the autozygous and non-autozygous portions of the genome using a t-test where the autozygous and non-autozygous rates are paired. To adjust for inconsistencies that might arise due to sampling error from individuals with small regions of autozygosity we further restrict our comparison to samples that have a total autozygous length of >=5%. We then examine the results of this statistic using different classes of variants **(fig. 2b).** We show as a control that rates of synonymous variants are not significantly reduced between the autozygous and non-autozygous sections, and then show a significant reduction in the rate at which we observe LOF variants within autozygous sections as compared to the non-autozygous sections, demonstrating direct selection against highly deleterious recessive variants.

###### S6.2 Quantifying the depletion of LOF genotypes

The previous analysis on variant genotypes within and outside of autozygous tracts suggests that synonymous variants are an effective neutral control, and that there has been selection against homozygous LOF variation. In the allele frequency range under 1% there were 16,163 segregating LOF sites, of which 847 were found as homozygotes. We then matched the LOF sites to randomly selected synonymous sites with the same frequency and observed how many of these were seen as homozygotes, repeating the random selection process 10,000 times to estimate the distribution of the number of LOF homozygotes expected under neutrality. **fig. 2A** shows the resulting distributions and LOF site counts. We see a 13.6% deficit in rare homozygous LOF sites compared to the mean of the distribution for matched synonymous sites. This estimate makes no use of the autozygosity assignment from **SM S3,** so is not biased by any inaccuracy in autozygosity assignment. It is an average over the range of allele frequencies below 1% homozygosed in our sample.

##### SM S7 The average number of recessive lethal variants carried by humans

The concept of lethal equivalents, defined as the expected number of heterozygous mutations in a single individual that would result in lethality when homozygosed, was first introduced in 1956, when it was estimated by a regression of the degrees of parental relatedness on the viabilities of their offspring(*46*). Whist methodologically sound, the estimate of the inbreeding coefficient of the infants was obtained theoretically by simply examining the known relationship of the parents. As we have seen, information obtained from recent pedigrees often does not capture the total amount of relatedness present in samples with extensive historic parental relatedness. Secondly, due to small sample sizes as well as the use of a theoretical inbreeding coefficient F, the results of this approach vary depending on the choice of the regression model used. Finally, the approach assumes that the socio-economic backgrounds as well as care received with regard to complications that might arise during pregnancy is the same across groups with different relatedness structures. From long standing cohort studies in the UK(*47*) we know that communities with more historic cousin marriage practices also have higher levels of health deprivation and education, which may have a significant impact on the birth outcome.

To minimize these limitations, we carried out a modern version of this approach by examining a dataset of 13776 mothers from the BiB cohort for which we had pregnancy outcomes. This dataset is particularly suited to this approach due to its large sample size from a single long-standing study in Bradford, UK. All of the mothers presented to the same maternity ward during their pregnancies and received standardised information on pre- and post-natal care. However the major difference from previous studies using this approach is that we had a direct DNA based estimate of the distribution of autozygosity (which we used in place of the the inbreeding coefficient) as a function of self-stated parental relatedness in this community from **fig S4A.** We chose to remove pregnancies with the birth of more than one child to further remove bias.

We calculated the Survivability (S) as the fraction of pregnancies that resulted in a healthy offspring surviving to at least one year of human life and carried out a weighted least squares regression as first described by Morton et al.(*46*) to estimate the coefficients and standard errors of the A and B terms in their model. In **fig 2C,** we report our estimates, as well as those obtained from other previously reported studies(*6*, *7*). The results obtained for A and B are 0.004970 (SE 0.001135, P=0.02205) and 0.22711 (SE 0.03745, P=0.00902) respectively. So as to be conservative with our results using this approach across all datasets, we choose to report the estimate of the number of recessive variants incompatible with healthy life in a human genome as B±SE.

Along with these estimates based on epidemiological data, we also obtain a direct estimate of the number of recessive lethal variants based on the suppression of homozygote genotypes **(SM S6.2).** For each allele frequency, we use the method of **SM S6.2** to calculate the number of variants that are depleted compared to the neutral expectation of synonymous variants. We then take a weighted sum with the allele frequency to get the total number of LOF variants (not sites) that are expected to be incompatible with adult life across 3,222 individuals. Standard errors are computed by a block jack-knife across individuals. As these variants are found in all 3,222 individuals we take the average to obtain an estimate for the number of heterozygous variants carried by a single individual that would be lethal or result in severe disease if homozygosed. In **fig. 2C,** we also include another recent direct estimate for a similar measure using recessive disease pedigrees(*48*).

##### SM S8 Analysis of LOF load in different population cohorts and relation to demographic structure

We carried out a comparative analysis of variants at functional loci and examined the mutational burden of LOF variants as identified by annotation between populations. To obtain a comparative dataset to use for the analysis, we obtained calls for worldwide populations from the 1000 Genomes Project phase 3 and restricted analysis to the same exome bait regions as our dataset and ran our annotation pipeline using the same settings **(SM S2).** We then annotated ancestral and derived alleles using Ensembl Compara’s 8 primates EPO alignment (http://www.ensembl.org/info/genome/compara/).

We calculated a statistic described in (8, 44) which compares two populations, given a particular category of sites, in terms of the number of derived alleles found at sites within that category in one population rather than the other. The rationale behind the statistic is to compare the haploid load of mutations of a particular class in one population versus another. If the mutation rate per year is identical in both populations after the two populations have diverged and have undergone different demographic patterns, then the difference in the haploid mutational load has to have occurred due to selection. To aggregate information across multiple individuals, we use observed derived allele frequencies in each population and compute the statistic as follows. At each site *i* we write the observed derived allele frequency in population A as 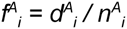, where *n^A^*, is the total number of alleles called there in population A and 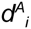, is the number of derived alleles called. Similarly we define Z^8^, in population B. Then if C is a particular category of protein-coding sites and S a set of synonymous sites, we define

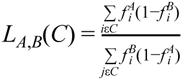

as a measure of the relative number of derived alleles found more often in population A compared to population B. We then define the ratio

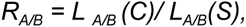

normalising by the value over putatively neutral synonymous sites, to mitigate any population-specific differences in overall mutation rate, as well as population-specific reference biases and any calling biases between our dataset and those from the 1000 Genomes. This is akin to the approaches in (*8*, *44*) where biases to do with branch shortening and deamination errors between ancient and modern genomes are mitigated. Estimates of the variance in *R_A/B_* were obtained using 100 block jackknifes on the set of sites in *C*.

Using our definition of LOF variants we then compared the value of *R_A/B_* in the 1000 Genomes Project cohort and our dataset to look for differences in historic selection for variants of this specific class. **Fig. 2D** shows R*_A/B_* for LOF variants found in one population as a comparison with that from an outbred European ancestry population, CEU. In most populations there is no significant difference. However, in the Finnish population sample (FIN), which underwent a severe bottleneck, we see significant differences compared to many other populations **(Table S6).** Our population (BB in **Fig. 2D),** which unlike FIN has a high heterozygosity so shows no evidence of a comparable bottleneck, shows a similar reduction in to FIN. We conclude that the reduction in load of severely deleterious mutations caused by homozygosing arising from endogamy in the BB population has been comparable to that arising from increased homozygosity during the bottleneck in the FIN population. We note that the CHS population from rural Southern China also has an increased R_A/B_ value without a decrease in heterozygosity, indicating that it may also have been subject to historical endogamy. The relatively high value of R^ for the GBR population may be a consequence of one third of its samples coming from Orkney, a small island archipelago north of Scotland with a small long term endogamous population.

##### SM S9 Comparative genomics, human versus mouse

In order to understand the accuracy with which mouse models reflect human phenotype, we compared 215 genes with rhLOF in our dataset to mouse gene knockout data, requiring an exact 1:1 mouse:human gene ortholog. We removed genes with i) a one to many, or a many to one cross-species mapping; ii) removed genes where the human ortholog also has an OMIM annotation; iii) considered a mouse gene essential if in any strain or experiment reported in the Jackson Labs Mouse Genome Informatics Mammalian Phenotype database that there was a lethal phenotype observed. Of these, there were 52 genes where a lethal mouse phenotype had been reported on at least one genetic background. Properties of genes essential in mouse but not in humans showed no significant differences to those non-essential in both species across protein divergence (dN/dS), number of gene duplications in humans since species divergence, and gene expression **(Fig S7)** (all data downloaded from Ensembl BioMart).

##### SM S10 Sanger sequencing validation

We selected 19 subjects for validation: 12 because of a rhLOF genotype in a highly expressed blood gene (for use in RNA expression in blood studies), and 7 with diverse genes of potential interest. Of the 37 homozygous LOF genotypes in these subjects, we validated 35 rhLOF genotypes (2 could not be assayed) using a different method (Sanger dideoxy sequencing) in these independent recall samples.

Sanger sequencing was performed on PCR products using an ABI 3730xl DNA analyser and ABI big dye terminator 3.1 cycle chemistry. We sequenced all samples with rare variant allele genotypes, and a control sample, for the sites selected.

##### SM S11 Western blot and RNA expression validation

Selected subjects were recalled for a further blood sample **(SM S10).** Whole blood was preserved for RNA immediately upon venesection using PAXgene system (Qiagen). Heparinized whole blood for protein assays was stored/transported at room temperature overnight and peripheral blood mononuclear cells were isolated by density gradient centrifugation with Lymphoprep (Stemcell Technologies).

Peripheral blood mononuclear cells were lysed using CelLytic M plus protease inhibitor (Sigma). Cell extracts were quantified by BCA (Pierce) and stored at -80°C till required. 2–10µg protein/lane were run on either 4–12% Tris Glycine or 10–20% Tricine Novex pre-cast gels (Life Technologies). Gel type used was determined by predicted protein size, specifically DPYD Tris-Gly gel 2ug; GCA Tricine gel 7.5ug; LSP1 Tris-Gly gel 3ug; SAMD9 Tris-Gly gel 10ug and MSRA Tricine gel 10ug. Gel electrophoresis and transfer to Novex 0.45μm PVDF membranes (Life technologies) were done using the XCell surelock system (Life Technologies). Membranes were incubated with antibody at 4°C overnight before development using Pierce Fast Western SuperSignal West Pico kit (Fisher) and imaged using the hyperprocessor (Amersham Pharmacia Biotech) with CL-Xposure Film (Life Technologies). Membranes were stripped with Restore (Thermo Scientific) and reprobed with antibody against α-tubulin (α-tub, Abeam) as loading control.

RNA was extracted using PAXgene Blood RNA kit (Qiagen), quantified by NanoDrop 8000 UV-Vis Spectrophotometer (Thermo-Scientific) and Bioanalyser (Agilent). 250ng total RNA per sample was labelled with the Illumina total prep RNA amplification kit (Ambion), and 750ng of labelled sample then hybridised to Illumina HumanHT-12v4 Expression BeadChip according to manufacturers instructions. Quantile-quantile normalisation and analysis was performed using Illumina GenomeStudio.

##### SM S12 Manual functional annotation of variants.

For each genotype we manually reviewed final bam sequences in the IGV viewer 2.3.59 (short read sequence data) for the rhLOF individual of interest, and 2 or more control subjects. We specifically looked for other nearby indels that would restore reading frame for rhLOF frameshift indels, and for adjacent variants that would eliminate stop codons, in addition to suspicious read alignment anomalies. We then examined the variant position in the UCSC genome browser, and in the ExAC browser, to look for potential issues with transcript and/or exon annotation that were missed by automated annotation with the LOFTEE tool. We reviewed rhLOF variants in annotated but non-conserved, alternative reading frames than the canonical exon. Splice variants in the last exon or 3’UTR, as well as ones with a nearby frame-restoring potential splice site, were also considered as suspect on the grounds that they were likely not to impact protein function.

##### SM S13 Primary healthcare records of Born In Bradford (BiB) subjects

All citizens of England are offered primary healthcare that is free at the point of use at general practices. Most practices have long used electronic health record (EHR) systems for the recording of diagnoses, symptoms, signs, according to the hierarchical system devised by Read (which maps to the internationally used SNOMED-CT) as well as prescriptions and test results. Structured primary care EHRs were obtained (and linked) for BiB participants registered with General Practitioner (GP) surgeries that use the TPP SystmOne platform. SystmOne has 100% coverage in Bradford and high coverage in surrounding areas. Records were extracted when the national unique health identifier (NHS number), surname, date of birth and gender were an exact match in SystmOne. From the full BiB cohort of 12450 mothers, 12333 (99.1%) were matched to their primary care records. Records were obtained from 18 months prior to study recruitment (Sep 2005 to Jun 2009), until end of November 2014, or until the participant died or withdrew from the cohort study if sooner. Total loss due to deaths, patients transferring to non-SystmOne practices, and withdrawing from the study, amounts to 0.4 years per person, with records available for 6.9 years per person, from a possible maximum 7.3 years per person. Of 2162 exome sequenced individuals available for analysis, 2145 (99.2%) had matched GP records.

For selected individuals with LOF variants of interest we also obtained lifetime electronic health records from SystmOne, and manually correlated Read codes with reported genotype-clinical phenotype associations in OMIM.

##### SM S14 Primary Healthcare record prescription- and consultation-rate analysis

Using the primary healthcare records of BiB subjects **(SM S13),** we performed a prescription rate analysis, assessing all classes of prescribed medicines, irrespective of indication. Drug prescriptions are captured in the primary care extract using 10-digit British National Formulary (BNF) IDs. Published data show a count of unique BNF chapter headings in a patient’s record can predict mortality and consultation rate(*20*). 10-digit BNF ID was truncated to a 4-digit chapter heading (the first two digits describe the organ system (e.g. cardiovascular), the second 2 digits the drug class (e.g. angiotensin converting enzyme inhibitors)), then distinct values per person were counted, so that multiple drug issues from the same BNF drug class were only counted once.

We calculated Consultation rate as clinical consultations per person year, using the formula 365 * (NCIinicalConsultationDays / DayslnSystem). A consultation event in SystmOne is an abstract event logged by the application when data is entered by a user. To avoid over-counting due to use of multiple SystmOne client sessions in one patient visit, days on which at least one consultation event occurred were counted (NCIinicalConsultationDays), with the following criteria: Include consultation events entered in a general practice setting; Include where the staff member role is GP/doctor or nurse; Include where consultation type is “Clinical”; Include where consultation method indicates face to face, telephone, home visit or unrecorded, but exclude others. The number of days per person transmitting GP data was derived from the SystmOne patient registration history. The extract begins 18 months prior to participant recruitment to the study. It ends (a) at end of November 2014, or (b) when the person withdraws from the study or (c) at death, whichever is earliest. A person can also transfer out of the system by registering with a non-SystmOne practice. These transfers out and back in were obtained from the SystmOne patient registration history, and the total number of days in the system per person was counted (DayslnSystem).

DayslnSystem was also used as a covariate in adjusted models, except where consultation rate was also in the model, as consultation rate already includes this component. Patient age is also used as a covariate in adjusted models, computed as age in years at end of last SystmOne GP registration period. Deprivation has been shown to predict GP consultation rate(*49*). Conventional measures of deprivation may lack validity across ethnic groups due to cultural differences in economic priorities and opportunities(*50*), whereas education has been shown to be effective in capturing variation in socio-economic position (SEP) across UK ethnic groups. The covariate education, with international qualifications equivalised based on UK NARIC [http://ecctis.co.uk/naric/ accessed 13 Feb 2015], was included in adjusted models as a marker for deprivation.

The effect of LOF genotypes on prescribing and consultation rate was examined by logistic regression using Stata v13. Patient age, education and days in the system were covariates for adjustment for all analyses. For consultation rate analysis, we also used a measure of mother’s education level as the best BiB individual-level marker for deprivation.

A small proportion of participants were in the system and providing data for a very short period that may not be sufficient to detect differences in disease burden. By way of sensitivity analysis, the above analyses were repeated only including patients who provided >5 person years of GP data (n=2020). Some authors have noted that individuals at the same level of education are not necessarily comparable in terms of socioeconomic (SES), and that multiple SES indicators should be used(*51*). Previous work in the BiB cohort has shown that SES variation within Pakistani ethnicity participants is captured by education, receipt of means tested benefits, material deprivation, subjective poverty and employment status(*52*). A third set of sensitivity analyses included these multiple SES indicators. None of the sensitivity analyses described here indicated conclusions that differed from the main analyses.

##### SM S15 Analysis of a subject with a predicted *PRDM9* knockout

The mechanism behind the localization and regulation of meiotic recombination in humans and other mammals are of considerable interest. Much research has been focussed on understanding the role of a single rapidly evolving gene, PR domain-containing 9 (*PRDM9*) in the molecular control of the distribution of meiotic double stranded breaks (DSBs) in mammals(*53*). Through the use of population based genome-wide analyses, bulk sperm sequencing as well as genome editing in model organisms, meiotic DSB sites have been characterised at high resolution.

Recent efforts have focused on understanding the patterns of recombination in species lacking PRDM9(*54*, *55*) as well as mouse *Prdm9* knockouts(*27*). These results suggest that in the absence of PRDM9 to localize breakpoints, most recombination is initiated at promoters and at other sites of PRDM9-independent H3K4 trimethylation. In humans, there has been extensive evidence to suggest that PRDM9 initiates and localises DSBs(*56*, *57*) and is a major determinant of hotspots. However, insights into its essentiality as well as direct functional studies in-vivo in humans have not been carried out. Here, we identified as part of our study an individual containing a homozygous knockout in the PRDM9 gene, within a long autozygous region. We also studied one of three children (others in the family did not consent to research). We performed further experiments to define the recombination landscape in this family.

###### SM S15.1 Validation of genotype and functional validation

We examined a pileup of the exome sequencing reads for the variant as well as standard Sanger dideoxy-sequencing validation of the variant to confirm the quality of the genotype call **(SM S10, fig. S6A,B).** We also observed **10** additional (not closely related) individuals in our study population to be heterozygous at this variant. Heterozygote variants (but no non-reference homozygotes) at this genotype were also observed in ExAC (http://exac.broadinstitute.org/variant/5–23524525-C-T). No homozygotes for any other *PRDM9* LOF variants were observed in ExAC. We carried out careful manual annotation of the variant to ensure that the variant would result in a homozygous knockout of the gene **(SM S12).**

###### SM S15.2 Cell line studies

No primary cell lines or immortalised human cell lines stably expressing *PRDM9* were available, as *PRDM9* is specifically expressed only at meiosis. We therefore performed site directed mutagenesis to generate the chr5:23524525 T allele using the QuikChange II mutagenesis kit (Agilent Technologies) and full-length *PRDM9* cDNA cloned into the pCEP4 expression plasmid as described(*58*), and confirmed the expected allele and insert by sequencing. We transfected the plasmid into HEK293 cell lines as described(*58*), and assayed protein by western blot using anti-FLAG M2 (Sigma).

###### SM S15.3 Illumina HiSeq X Ten whole genome sequencing

Whole genome sequencing data for the PRDM9 duo was carried out by extracting DNA as described in **SM 1.2.** A single library (650 base pair inserts) was constructed for each sample. The libraries were multiplexed and sequenced across several lanes on the Illumina HiSeq X platform (paired-end sequencing, 151 cycles). Variant calling and filtering were carried out using the approaches described in **SM 1.5.**

###### SM S15.4 10XGenomics long range Gemcode molecular phasing

The 10X Genomics GemCode reagent delivery system partitions long DNA molecules (including DNA >100 kb) and prepares sequencing libraries in parallel across the partitions such that all fragments produced within a partition share a common barcode. A simple workflow combines large partition numbers with a massively diverse barcode library to generate >100,000 barcode containing partitions. Libraries on the mother and child were sequenced on an Illumina HiSeq 2500 instrument in Rapid Run mode. The GemCode Long Ranger Software maps short read data to original long molecules using the barcodes provided by the reagent delivery system, thus providing long range phasing information.

The DNA samples available for the PRDM9 duo, whilst of adequately high molecular weight for most genomics applications, were more fragmented than for the Coriell reference samples NA12878 and NA12882 **(Table S7).** Nonetheless, >90% of SNPs could be phased in each of the four samples. Unlike, e.g. detection of structural variants, identification of recombination breakpoints (SM S15.5) is less sensitive to the length of phased segments.

###### SM S15.5 Obtaining recombination breakpoints using phased parent-child duos

We identify the recombination sites in the maternal meiosis leading to the child’s genome using heterozygous sites in the mother and child that are informative about the transmitted chromosome. Of the possible allelic combinations of the mother and child at each site **(Table S8)** 4 combinations are informative if only the mother is phased while 8 are informative if both the mother and child are phased. We first obtained a gold standard phased set of calls for the NA12878 duo by using the CEPH 13 member pedigree and only considered sites for which the phase and genotype information are consistent across all members of the pedigree, who had been previously sequenced (*59*). This approach yielded a set of 53 crossovers with a mean interval length of 40.5kb.

We then used this approach on the data obtained from our phased whole genome sequencing **(SM 15.4).** Due to the higher rates of genotype error rate and phasing error rate, we initially only used phase information from the mother to locate the crossovers and consider bi-allelic SNPs that are of high quality (10XGenomics phase quality scores at maximum). Once we located such intervals we filtered for genotype and phasing errors that might have occurred by removing crossover intervals that lie within 300kb of each other. To ensure accuracy, we carried out manual visualisation of the crossover locations in the 10XGenomics Loupe browser to ensure that there were at least 2 informative SNPs on each end of a crossover that were of extremely high quality. We then refined the crossover intervals using the child’s phasing where it was available and overlapped with the phase set of the mother (this was in >90% of cases in NA12878 duo and ~70% in the PRDM9 duo), as well as considering sites where the phase quality was lower but was nevertheless consistent with the phased genotypes in the other sample of the duo.

This approach resulted in 42 crossovers in NA12878 / NA12882 duo, with a mean crossover interval length of 16.2kb and 37 crossovers in the PRDM9 duo with a mean crossover length of 51.8 kb. As a validation of our method, we show that 40/42 (95%) of the NA12878 duo crossovers also overlap those from the pedigree based gold standard, signifying that we are able to identify crossovers in parent-child duos with high specificity.

###### SM S15.6 Statistical analysis to infer recombination landscapes

In order to compare the recombination landscape of the two samples, we utilized a method that has been previously described in several studies that have compared crossovers obtained from pedigree based studies with those from the fine scale recombination map(*22*, *60*). Here, we seek to examine the PRDM9 hotspot usage phenotype by comparing the crossover events that we call using long range molecular phasing with those obtained from high-resolution maps of meiotic DSBs in individual human genomes(*25*). These are tightly coupled with crossover locations observed from population sequencing and linkage disequilibrium (LD) based methods but provide information on the localisation of DNA binding by PRDM9 at much finer scale.

For additional confirmation and to compare our results with those from these previous studies we also utilized 32,996 autosomal hotspot locations inferred from genome-wide Phase II HapMap LD data(*57*).

The method estimates the proportion of crossovers (*α*) that occur in a set of predefined feature intervals. Care must be taken because we do not know the precise location of the crossover, only an interval (possibly large) within which it took place. We utilise the previously published method almost exactly as referenced above but describe it again here due to slight changes in implementation. We begin by obtaining the probability that crossover r overlaps a feature:

P(r overlaps a feature) = *α +* (1-*α*)P(r overlaps a feature by chance)

We estimated P(r overlaps a feature by chance) by randomly shifting the crossover r a normally-distributed distance (mean 0, std. deviation 200kb) 100 times and counting the fraction of these moves that result in the crossover r overlapping a feature (note that this probability differs for each crossover, due to the size of the interval containing r and the density of features in the local region). The likelihood of *α* for a crossover r is *δ*_r_ P(r overlaps a feature) + (1-*δ*_r_)(1-P(r overlaps a feature)), where *δ*_r_ is an indicator variable that is 1 if r overlaps a feature and 0 if not. The likelihood of *α* for the full set of crossovers is the product of the per-crossover likelihoods.

As in the previous references, the likelihood of *α* for all crossovers localised to an interval smaller than 30kb was calculated over an interval of values between 0 and 1 in 100 steps and implemented using the NLM package in r. The 95% confidence interval of *α* was considered to include all values of *α* for which the log likelihood was within 2 units (the asymptotic cutoff) of the maximum log likelihood. Note that this estimation procedure naturally accounts for uncertainty in the location of LD hotspots and in the location of crossovers, as the overlap by chance is influenced by both the width of the estimated hotspots and the interval sizes in which we infer crossovers.

We report in the main text point numbers for the 23 crossovers in the PRDM9- duo and 34 crossovers in the NA12878 duo localised to within 30kb. When using all 37 (respectively 42) crossovers we get consistent results: for the PRDM9 A Union DSB hotspots: NA12878 duo - 55.2% [38%-71%], PRDM9- duo - 9.7% [0%-26%]; and for the LD based hotspots NA12878 - 72.1% [54%-86%], PRDM9--18.3% [2%-34%],

###### SM S15.7 Examination of crossover overlap with GC rich regions, promoters or H3k4me3 sites

We obtained GC content, gene promoter and promoter flanking region information from Ensembl(*43*), and consolidated imputed data for narrow contiguous regions of enrichment (peaks) for H3K4Me3 ChIP-seq data for multiple cell types from the epigenomics roadmap project(61). Using the same approach used in **SM15.5** for overlap of our regions with PRDM9 hotspots we found no significant differences between the proportion of crossovers overlapping H3K4Me3 sites (NA12878 duo 4% [0–8%], PRDM9 duo 5% [0–8%]), gene promoters (NA12878 duo 0% [0–4%], PRDM9 duo 0%[0–4%]) and their flanking regions, or in the GC content of 200kb of DNA (t.test p-value 0.643) from the middle position of each crossover interval.

##### SM S16 Druggability and clinical approval analysis

We annotated genes with LOF variants with information concerning potential druggability - that is the potential for modulation of the protein target by a water-soluble small molecule drug. Druggable proteins usually contain a defined binding pocket or active site, which could act as a site of action (pharmacophore) for an orally bioavailable small molecule drug. We grouped proteins into four druggability classes, based on a collation of complementary published annotations of the potentially druggable genome and publically available databases of small molecules in the drug gene interaction database ((DGIDB); http://daidb.aenome.wustl.edu/; (v1.72). Targets in class D1 have a known drug recorded in dgidb; class D2 have small molecule tools recorded in CHEMBL (www.ebi.ac.uk/chembl) which may be in current development within pharmaceutical companies, and could be used as tools in animal and cellular models; class 3 are homologous to class 1 or class 2 targets described in several druggability publications collated in DGIDB; class 4 are predicted to contain a potentially druggable pharmacophore based on de novo structure-based druggability prediction using the dogsitescorer tool(*62*) (dogsite.zbh.uni-hamburg.de).

A protein can be considered a potential biopharmaceutical target if it is present in the cell membrane or extracellular space. Consequently we designated a protein as a biopharmaceutical target if it was reported to be extracellular or transmembrane in the Gene Ontology location category(*63*).

We compared the ultimate success or failure of drug development for targets contained within the different LOF gene datasets using a recently published dataset of drug development outcomes for 19,085 target-indication pairs(*30*)(citeline.com/products/pharmaprojects/). We used a Chi-squared test to evaluate differences in the ultimate EU/US approval rate for targets with observed LOF variants, compared to background information target-indication pair approval.

##### SM S17 Protein-protein interaction network analysis

Collections of genes with loss of function (LOF) and gain of function (GOF) variants **(table S4)** were compared against a genome wide background of molecular interactions derived from the STRING database (string-db.org/). Interactions were organised into seven categories, Binding, Reaction, Activation, Expression, Catalysis, Post Translational Modification and All Interactions. The distribution of interactions were observed to be non-normal in most cases, probably due to missing data. We therefore compared the distributions of interactions between gene collections using a non-parametric Kruskal-Wallis test to obtain a p-value (R, http://www.r-proiect.org/). Low medians were seen across several of the seven interaction groups, therefore 5% and 95% quantiles were also reported in addition to median values for each group.

**Fig. S1.**
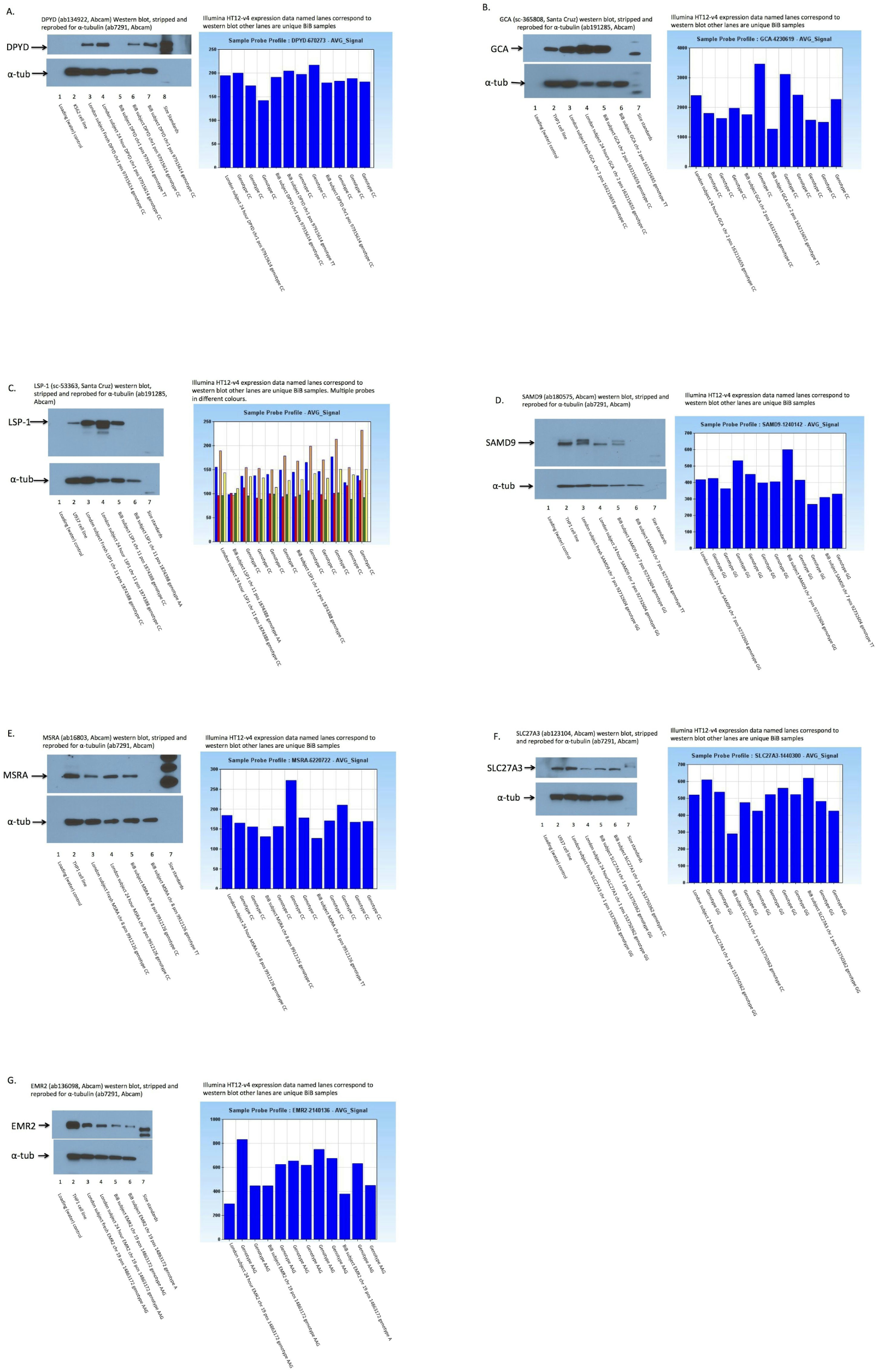
RNA and protein level validation of predicted LOF genotypes. Left panels show protein expression from Western blot. Right panels show probe-level Illumina HT-12v4 RNA expression array data. Panels A,B,C,D,E show absent protein in the LOF sample as assessed by Western blot. Panel G shows low protein in the LOF sample. Panel F shows apparently normal protein (presuming the antibody is specific) but low RNA levels in the LOF sample.

**Fig.S2.**
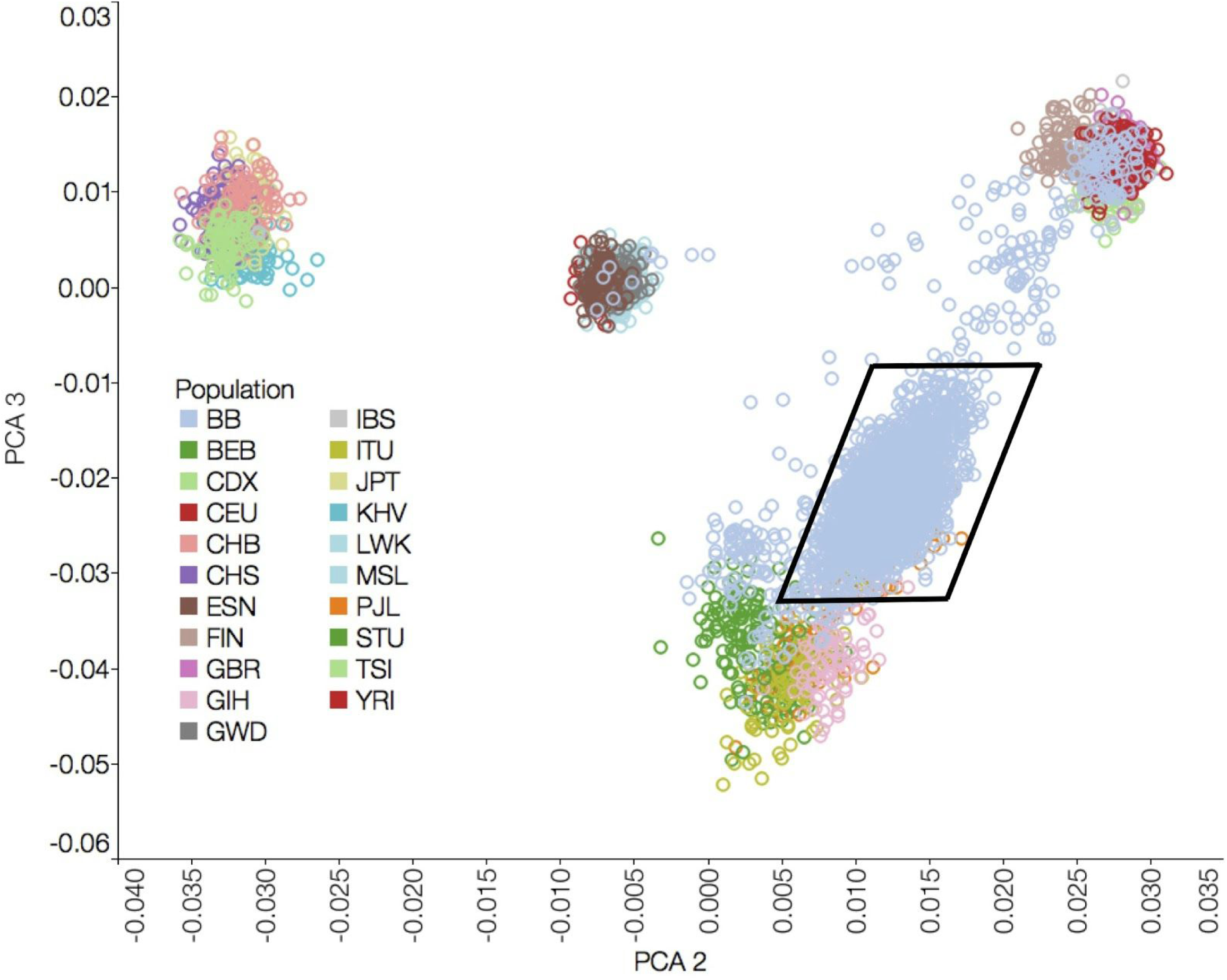
DNA-determined heritage. Principal component analysis was performed on individuals from Phase 3 of the 1000 Genomes Project, with the current dataset mapped onto this reference. We defined a polygon to include samples that were clustered together with the 1000 Genomes Project Pakistani population. Legends: BB - Birmingham and Born In Bradford; others -1000 Genomes Project ethnic groups. PCA polygon (in black) refers to the subset of samples retained in the BB dataset for further analysis

**Fig.S3.**
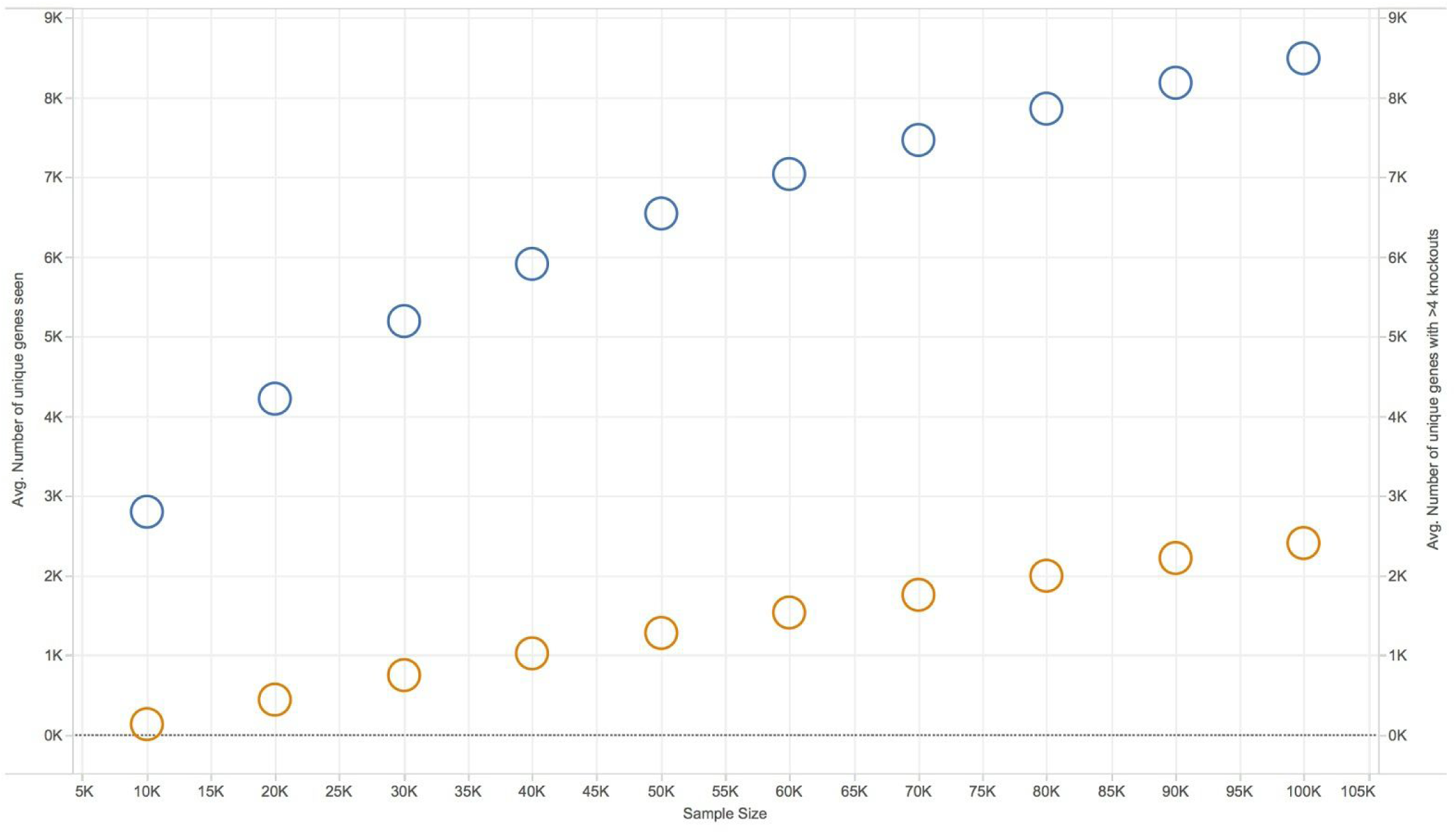
**Estimates of number of rare homozygous knockout genes seen in larger cohorts**. (blue circles), and number of rare homozygous knockout genes with more than 4 individuals (orange circles).

**Fig. S4.**
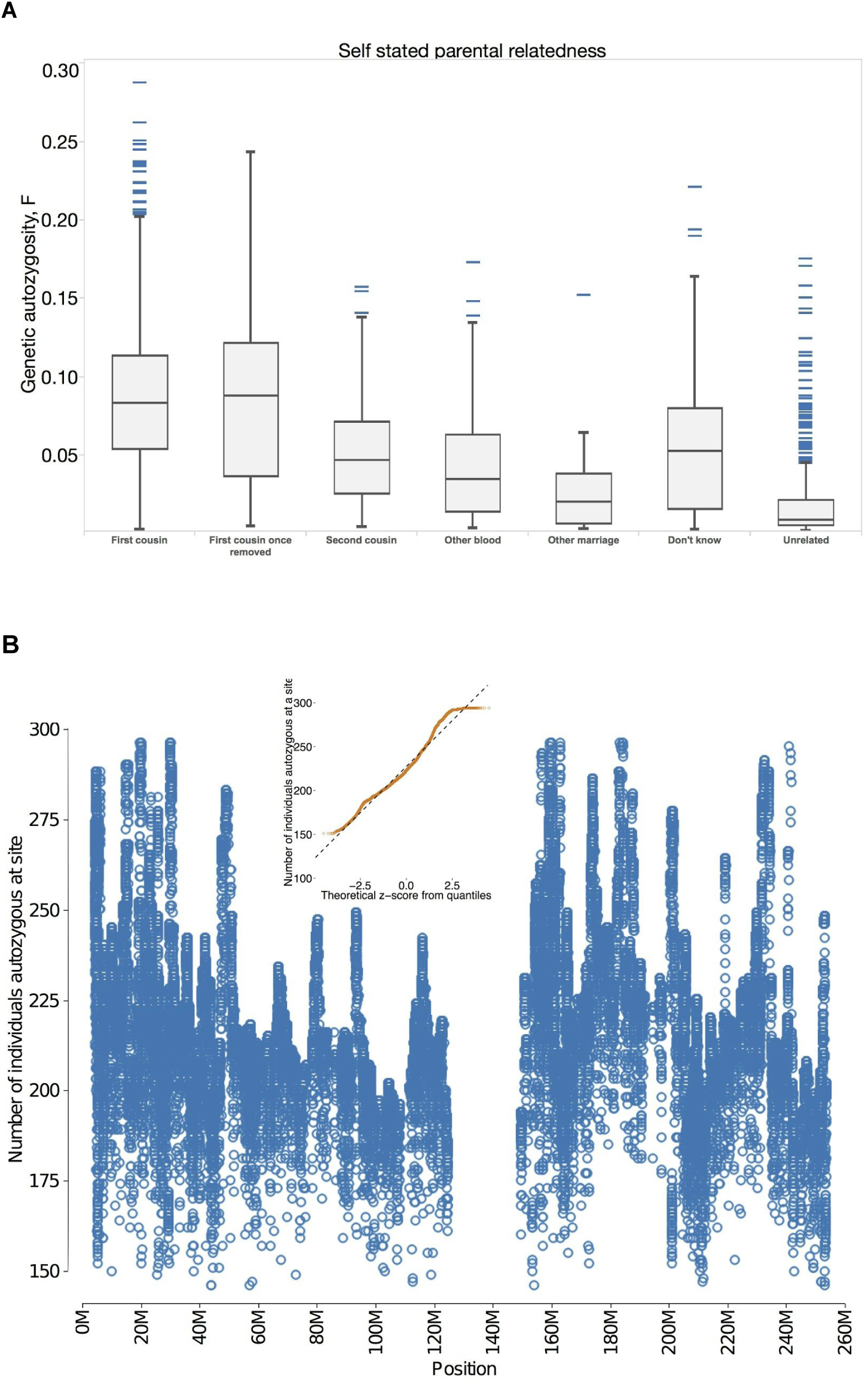
Analysis of autozygosity in populations. **(A)** Self-stated vs genetic autozygosity for individuals from Born In Bradford. Distribution of total length of genome in autozygous stretches shown in boxplots, with individuals for whom no data was available indicated in the ‘Did not answer’ column. Genetic autozygosity, declines with self-stated parental relatedness and reflects theoretical expectations. **(B)** Levels of autozygosity observed across the genome. Number of individuals autozygous at a site in the protein coding sections of the genome (blue circles) across positions on chromosome 1 (shown as a representative example) on the x-axis. Inset in orange circles is a Q-Q plot of the theoretical expectation of the distribution, and shows normality of the data.

**Fig. S5.**
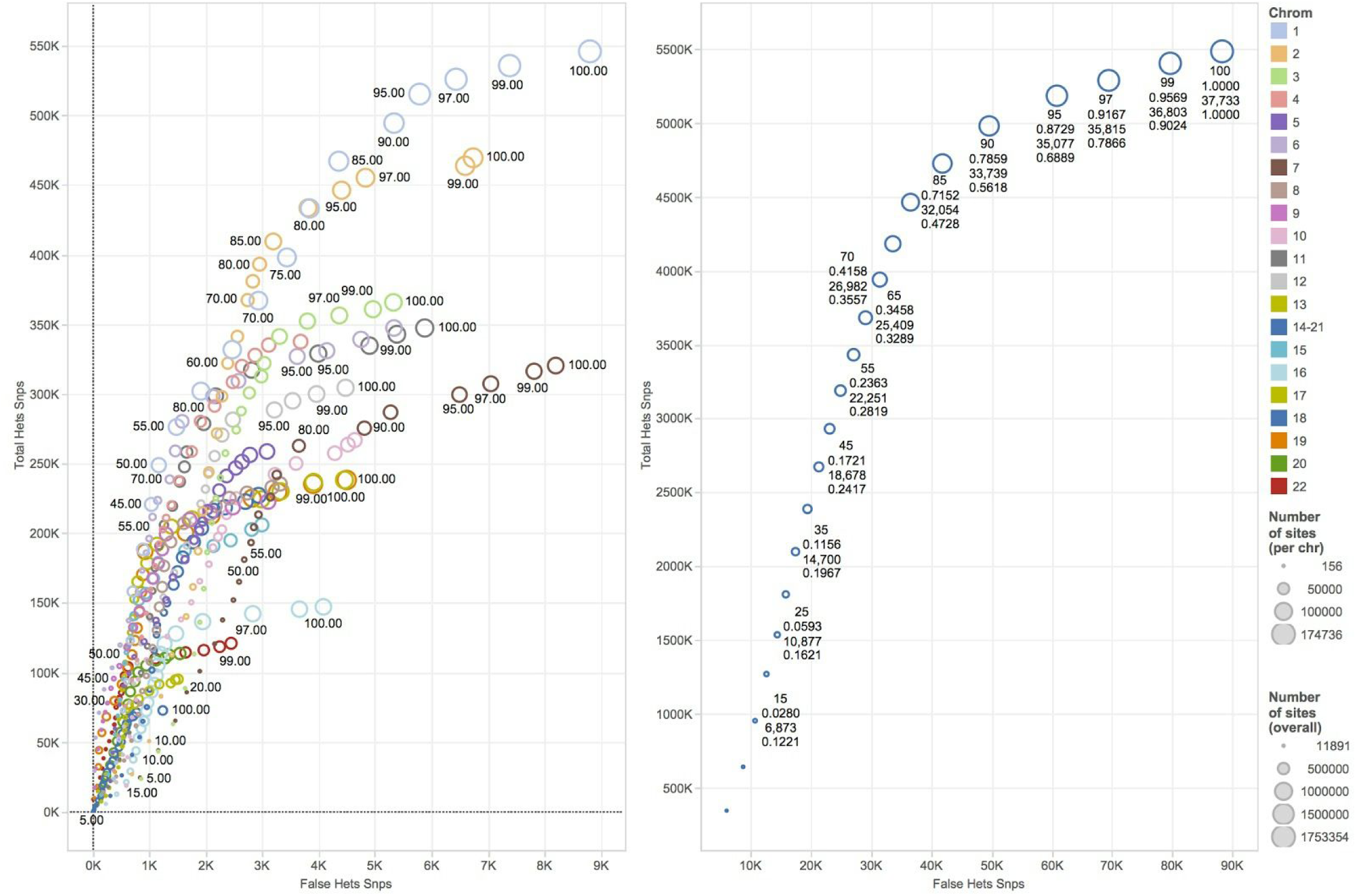
Using heterozygous calls within an autozygous section as a measure of variant quality control. Total number of discovered heterozygous sites in 3,222 individuals as a function of sites that were heterozygous within an autozygous block (x-axis) across individual chromosomes (colored circles on left panel) and summed across the genome (blue circles on right panel). Total number of sites overall is represented by the size of circle. Labels on each point represent in order: the VQSR false positive threshold reflecting the percentage of true positive sites left; (right panel only) the fraction of total sites left at that threshold, the mean number of hets per person and the number of false heterozygote sites left.

**Fig. S6.**
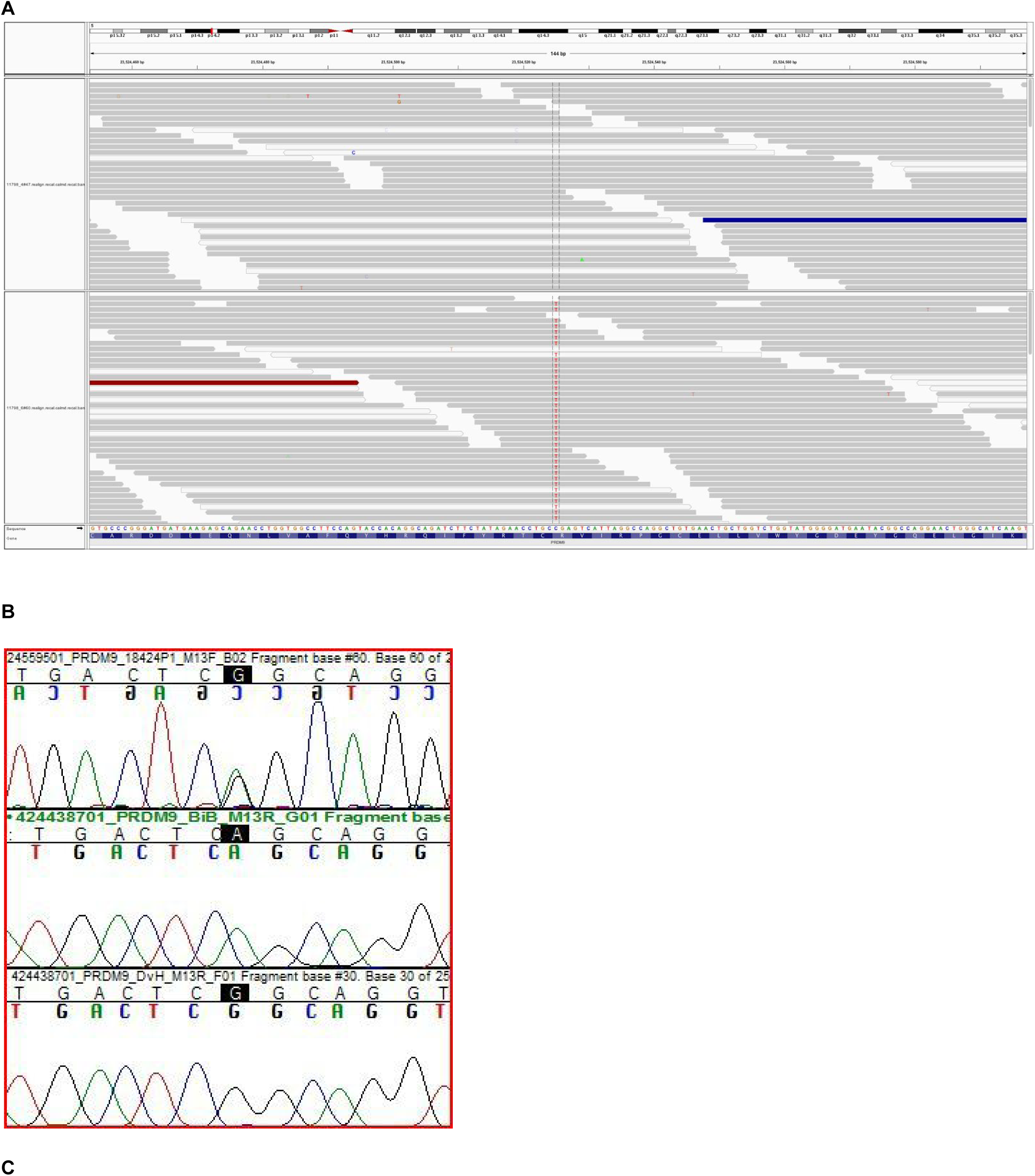
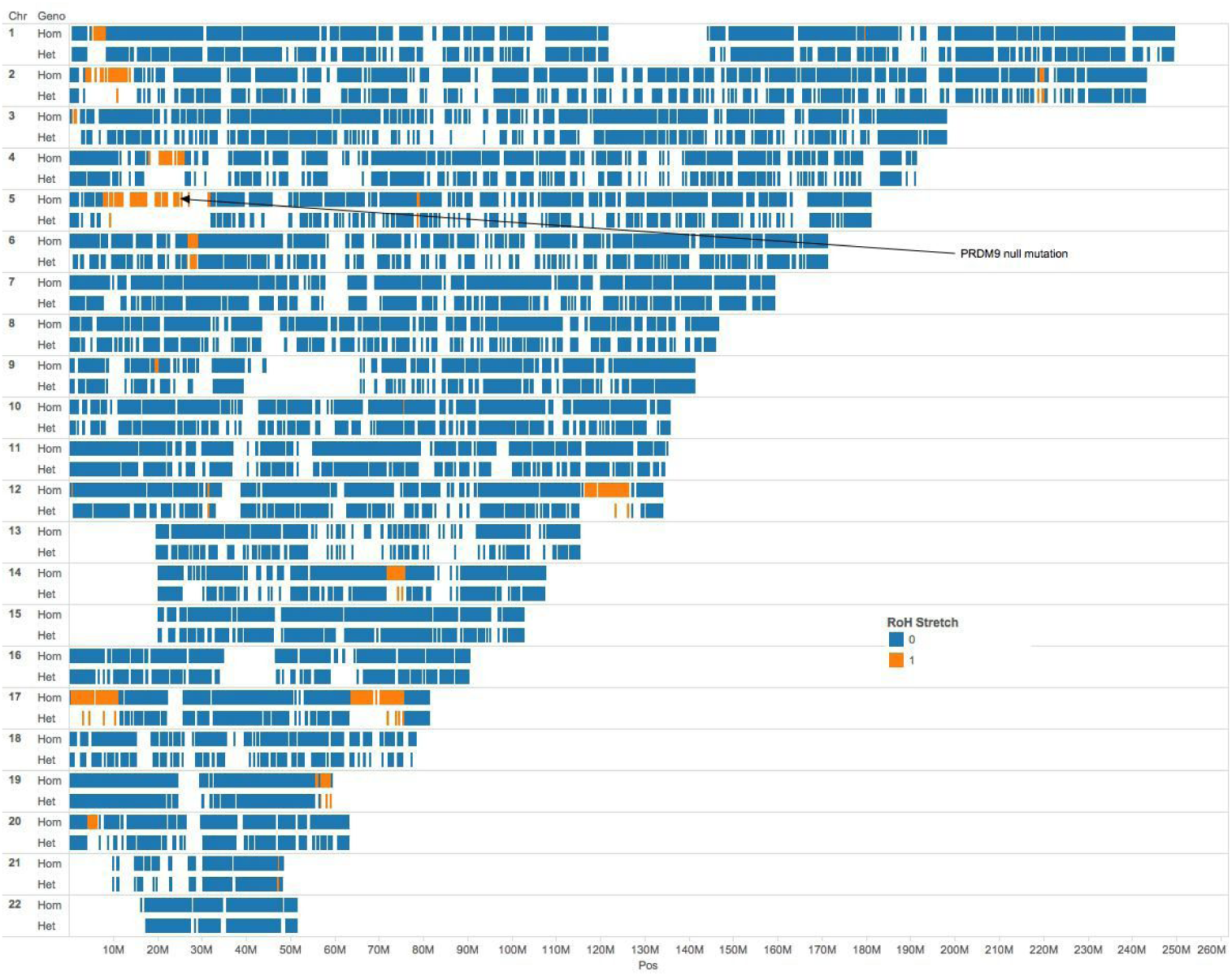
Genomic analysis of PRDM9 rhLOF mother and her child. **(A)** Exome sequencing reads showing PRDM9 chr5:23524525 C /T variant in a control (upper image, genotype CC) sample (upper), and in the mother (lower image, genotype TT). Images from the Integrated Genome Viewer **(SM S12). (B)** Sanger dideoxy sequence traces of child (upper trace). This confirms the expected heterozygous genotype and germline transmission of this allele. Middle trace mother, bottom trace control sample. (C) Genomic map of autozygous (identical-by-descent) regions in the mother, showing position of the PRDM9 rhLOF.

**Fig. S7.**
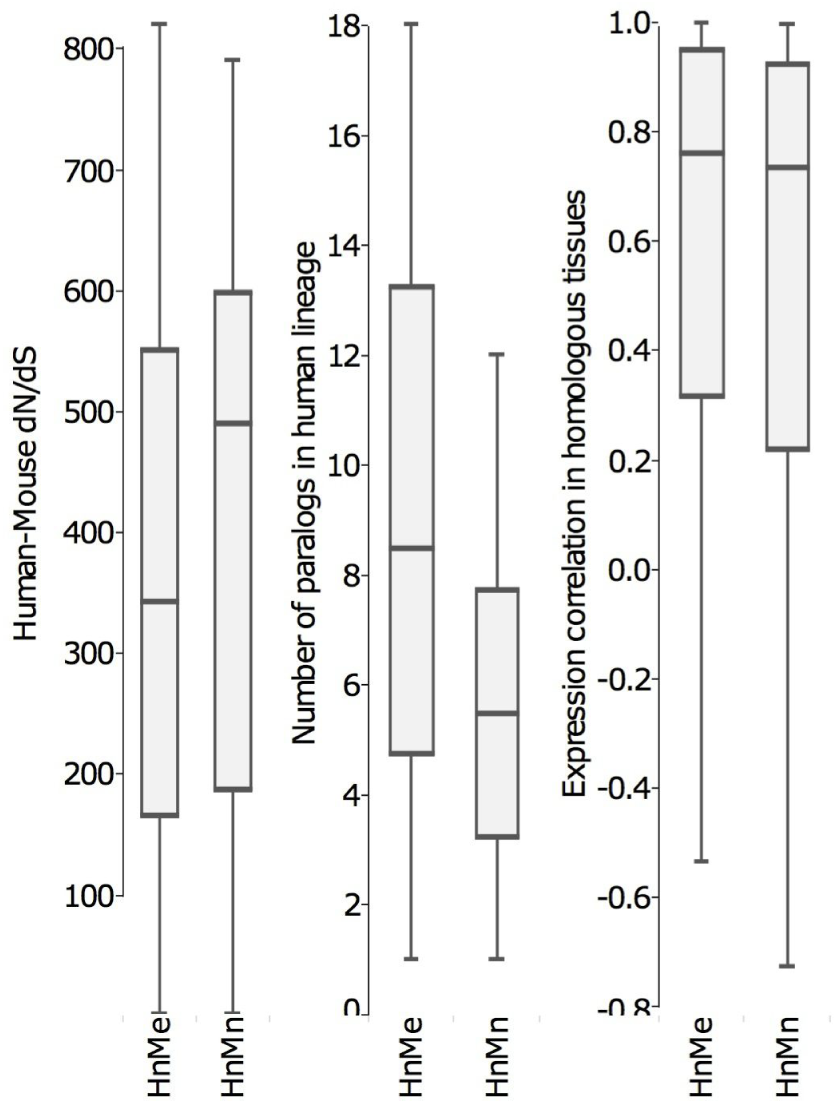
Mouse Human Orthologs. Properties of genes essential in mouse but not in humans (HnMe), compared with those non-essential in both species (HnMn) across protein sequence divergence (dN/dS), number of gene duplications in humans since species divergence and gene expression show no significant differences

### Supplementary Tables

**Table S1.**
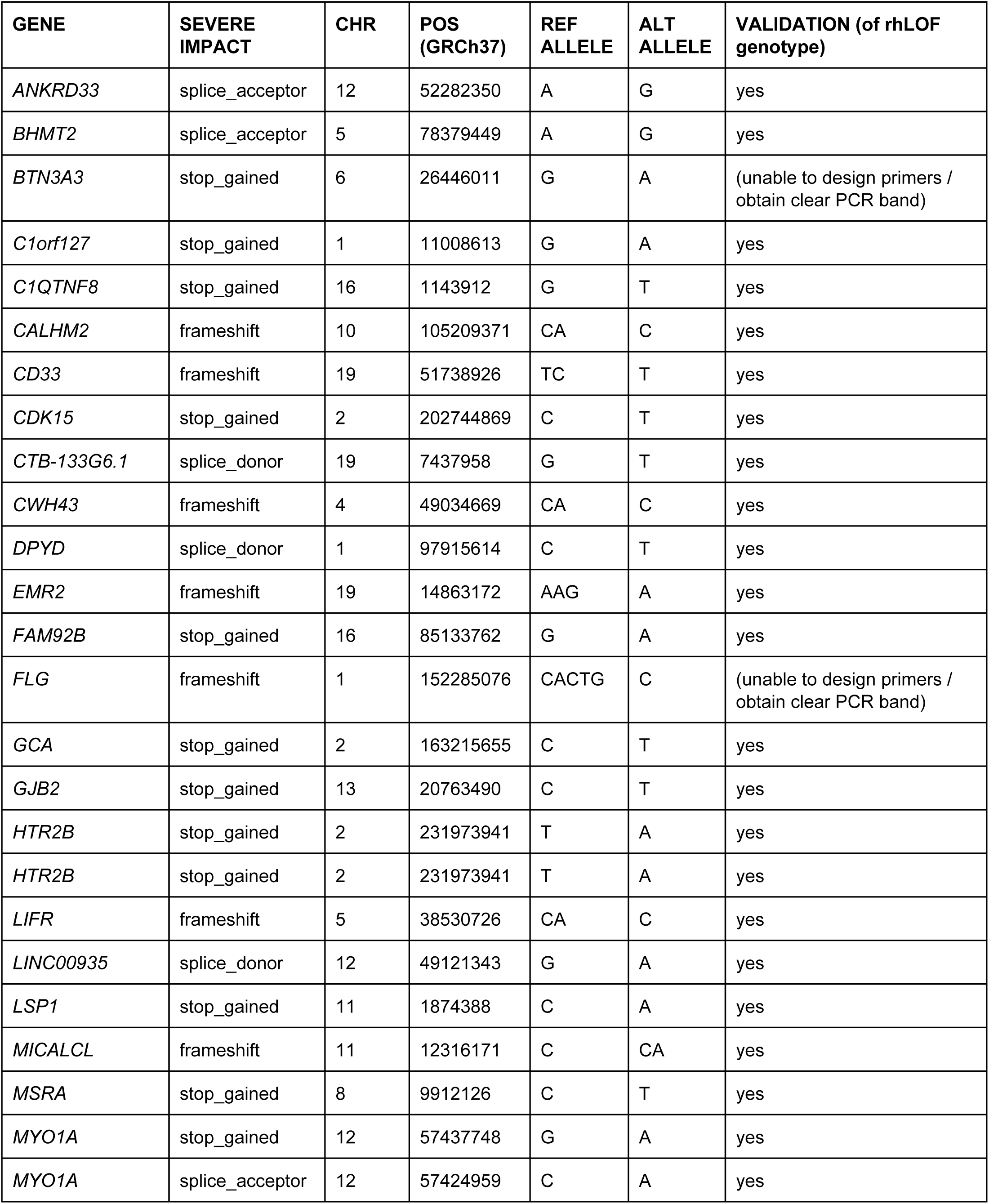
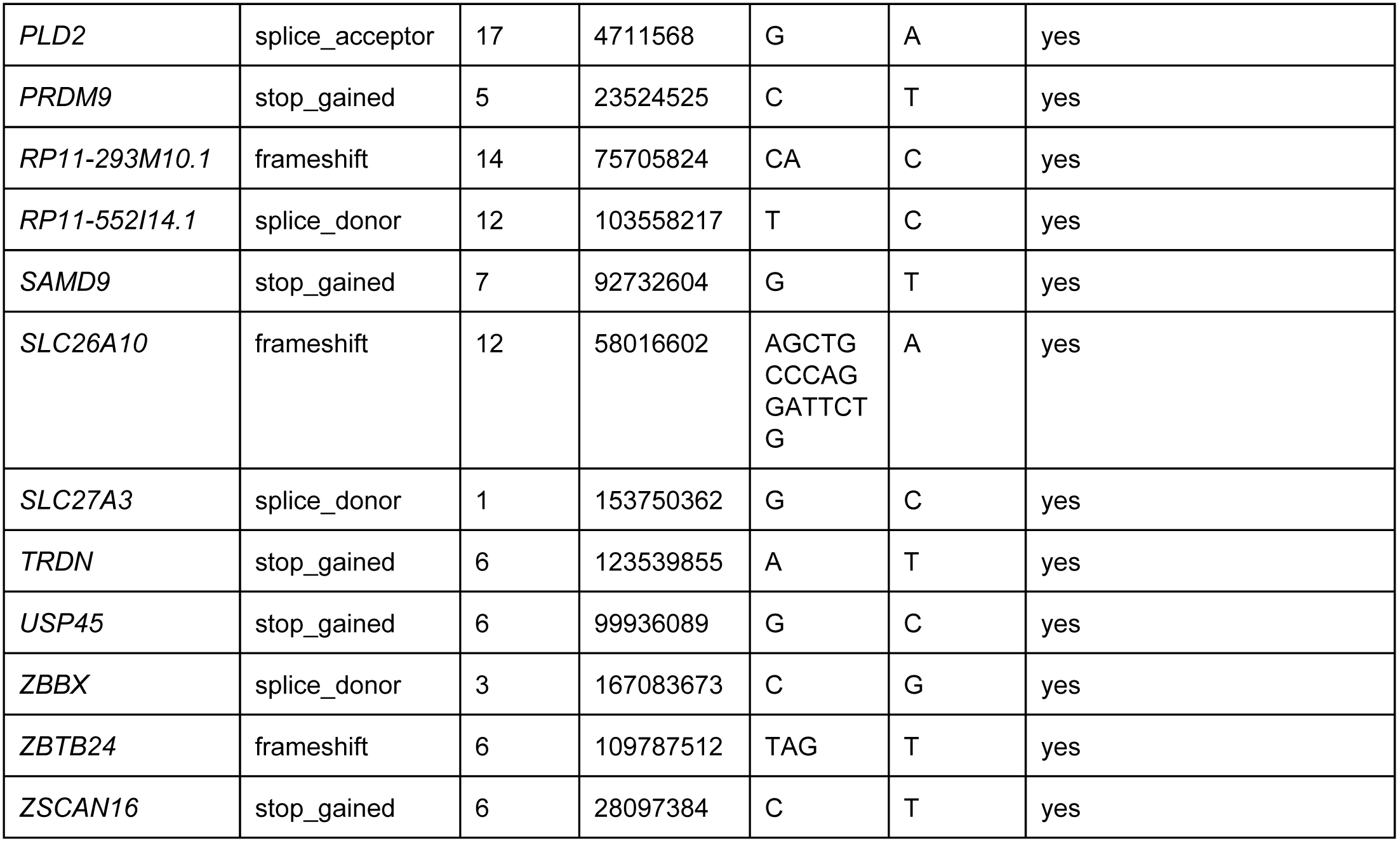
Sanger sequencing validation of variants identified by exome sequencing.

**Table S2.**
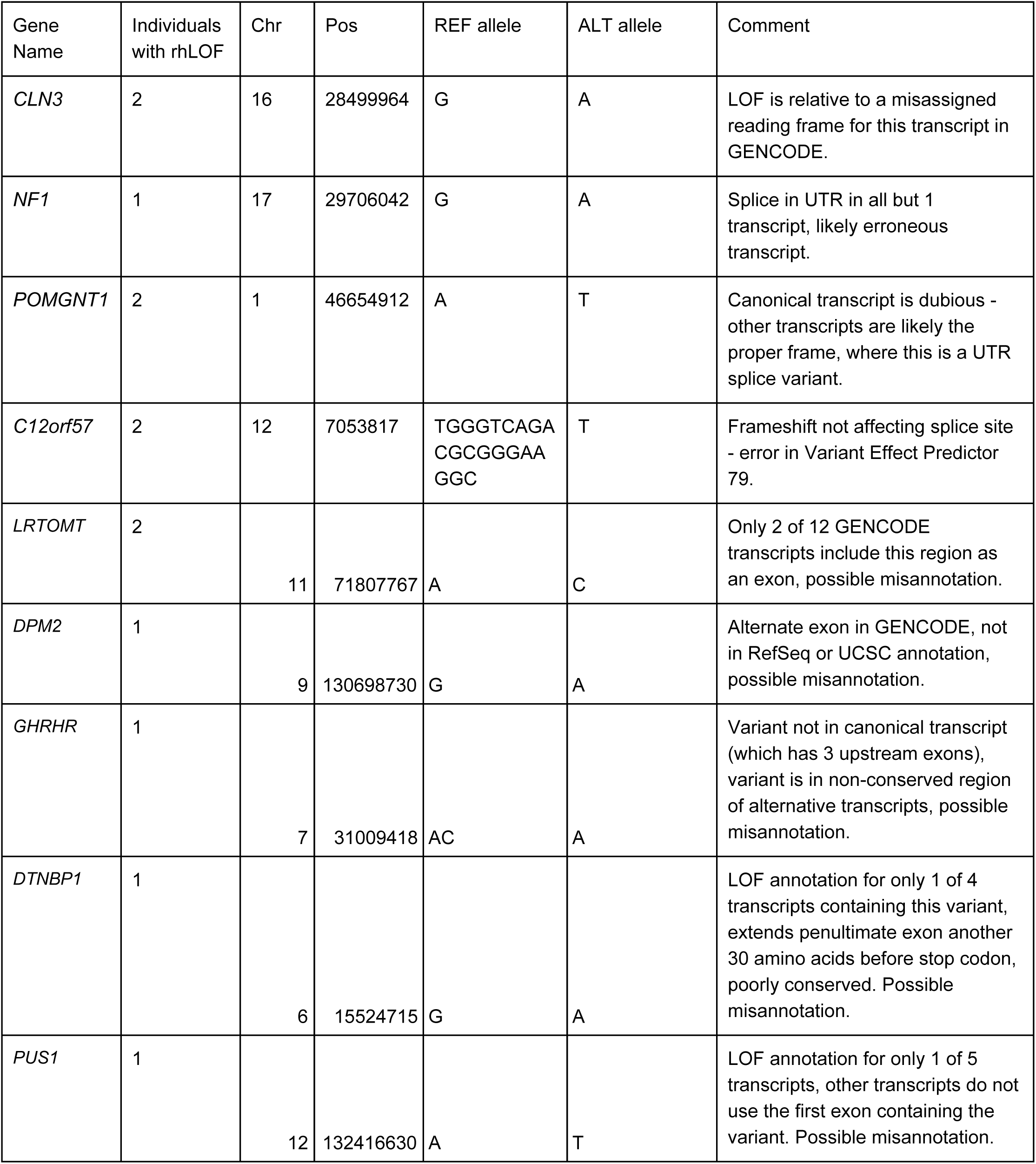
Variants in OMIM-disease genes erroneously annotated as LOF.

**Table S3.**
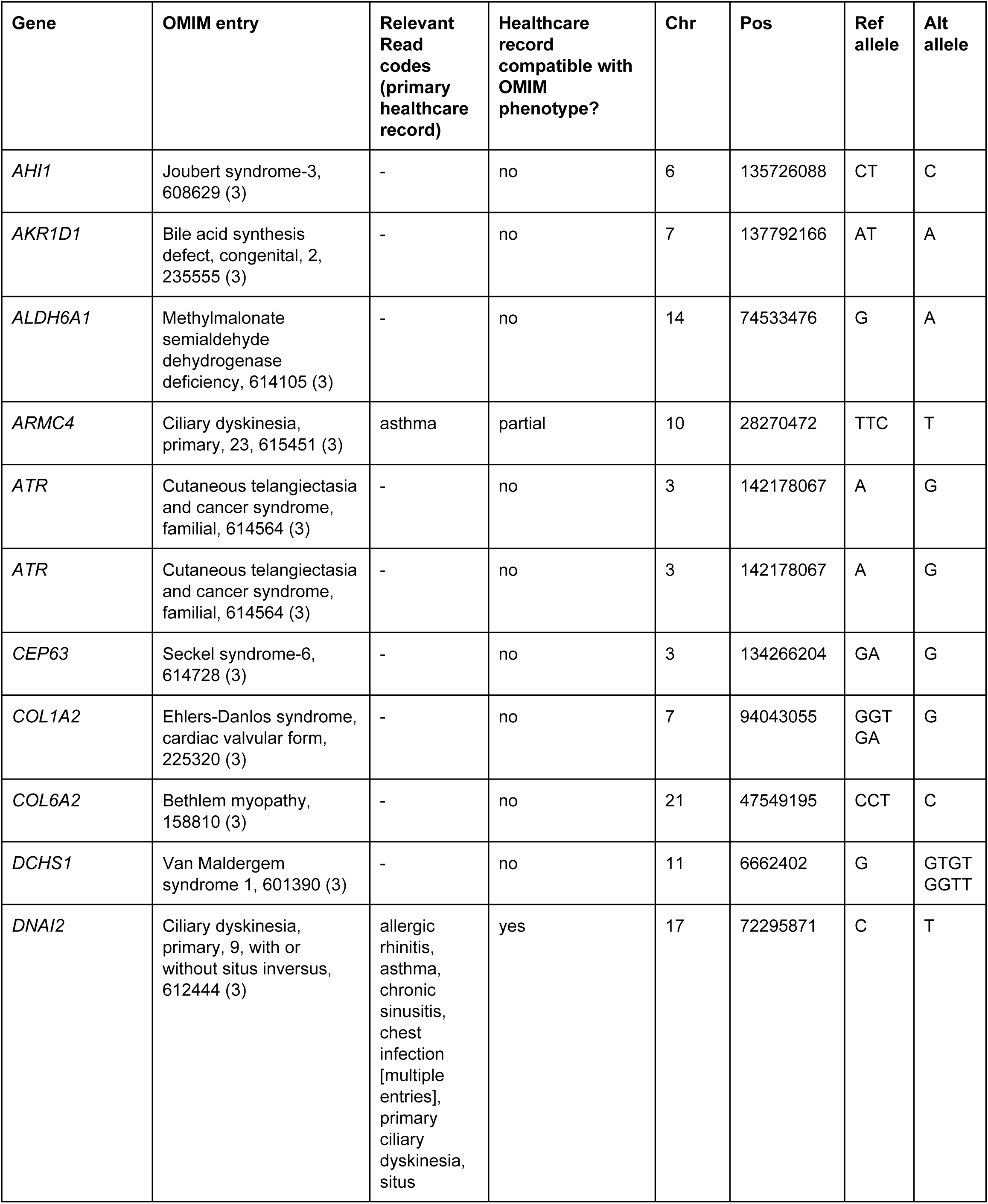
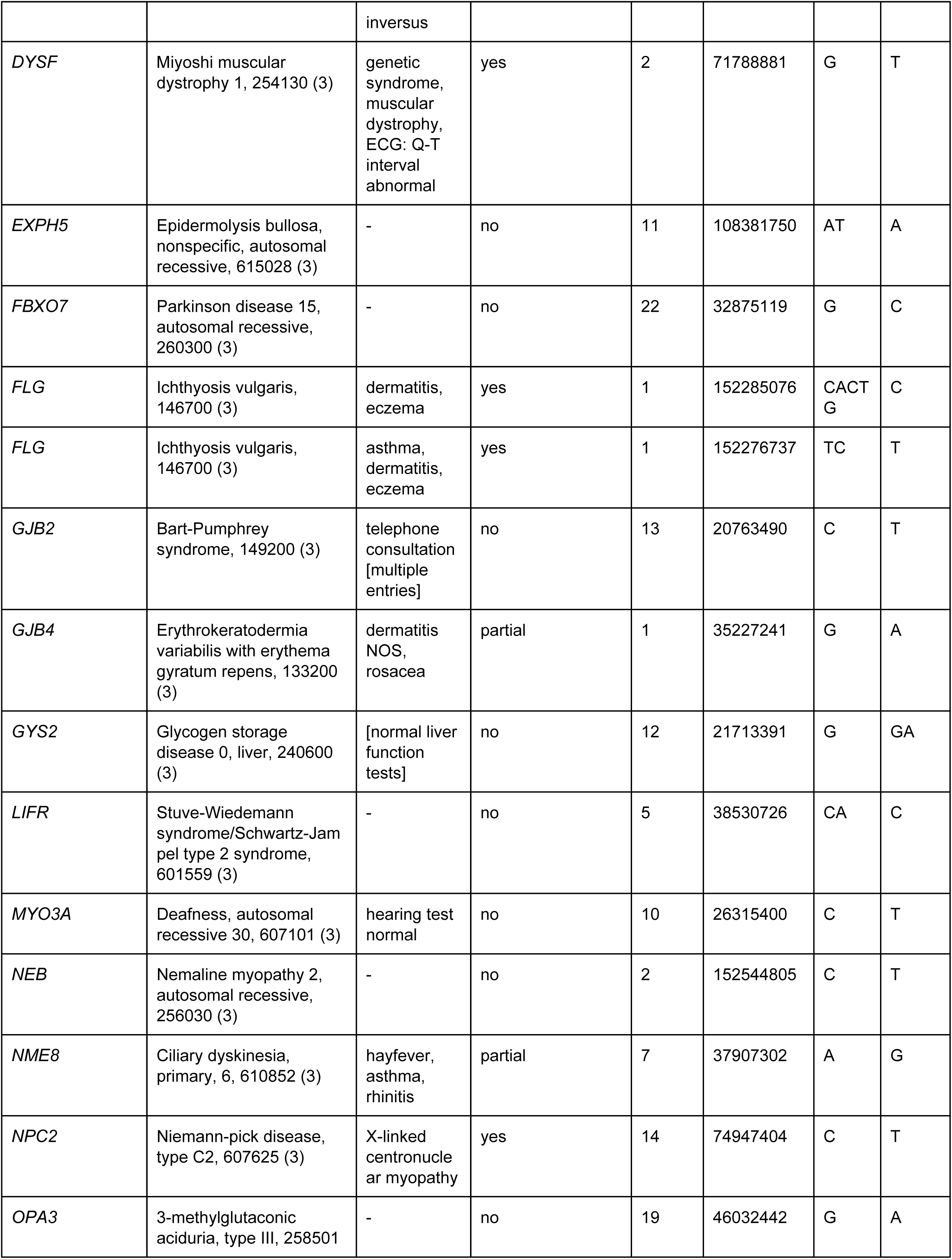
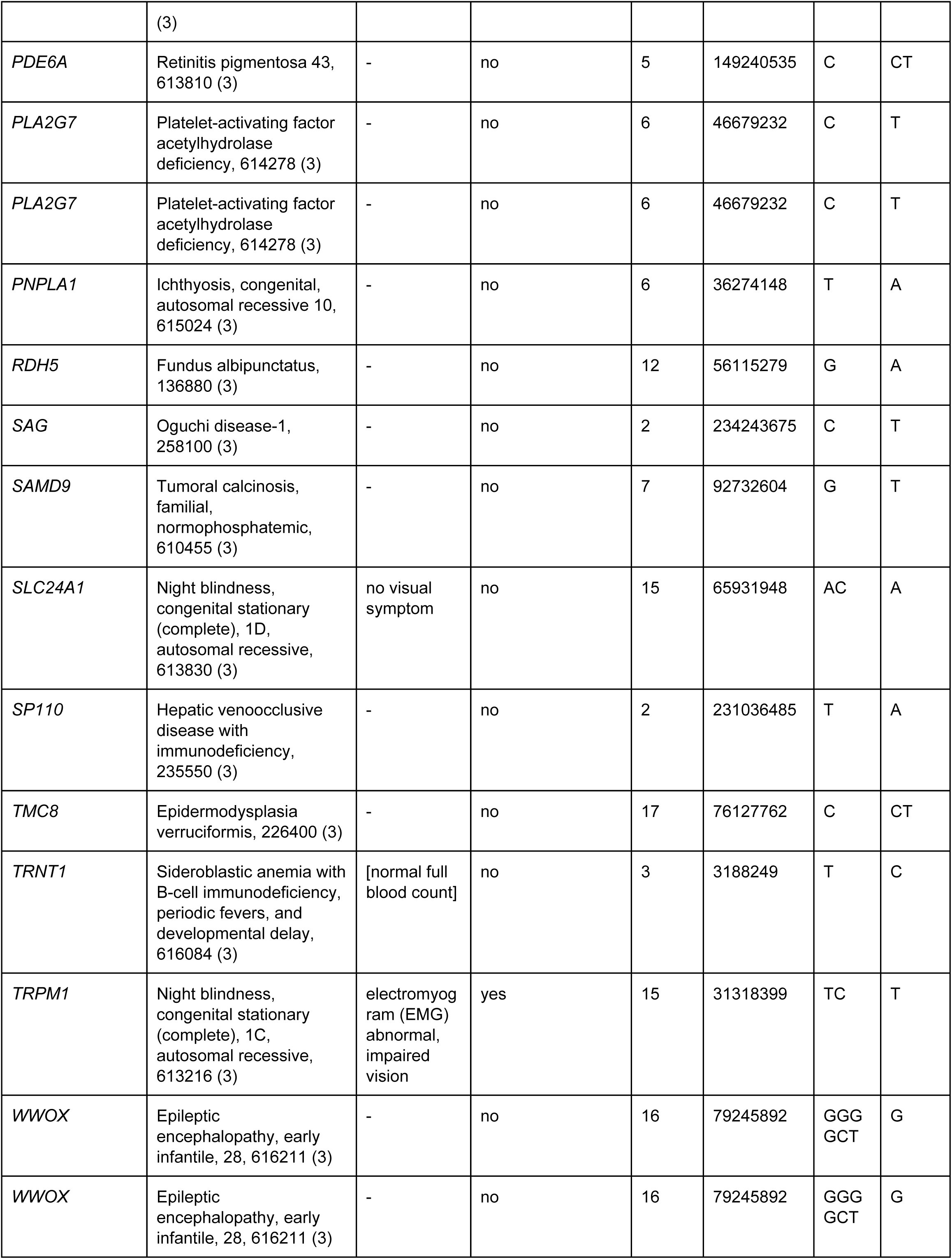
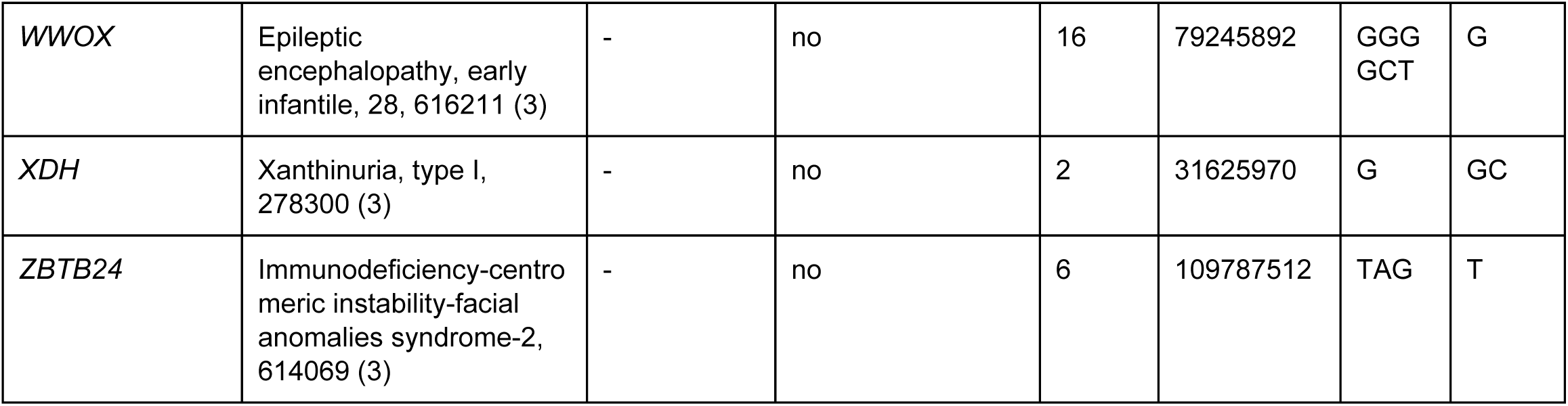
Presence in primary healthcare records of Born In Bradford subjects of phenotypes compatible with OMIM recessive genetic diseases. Autosomal recessive phenotypes only. Each row represents a sequenced individual. Entries marked [] are our interpretation/summary of healthcare record codes.

**Table S4.** A systematic evaluation of the degree of protein-protein interaction in genes carrying Loss of Function (LOF) variants. The following gene sets were compared against genome-wide interactions in the STRING dataset **(SM S17):** “Born in Bradford LOF all” (n=476), representing a non-redundant list of genes with observed LOF variants observed in this study in the Born in Bradford cohort. “Decode LOF all” (n=3769), represents all genes with LOF variants in an Icelandic sample(*3*), these genes are further divided into two subgroups “Decode LOF “benign”” (n=3530), a subset of genes with LOF variants identified in healthy subjects and “Decode LOF “pathogenic”” (n=416), a subset of genes with LOF variants identified in subjects with 1 or more offspring that died before age 15. The final two gene sets, “Orphanet GOF all” (n=106) and “Orphanet LOF all” (n=458), represent genes with pathogenic Mendelian gain of function and loss of function variants respectively reported in the Orphanet rare disease catalogue (www.orpha.net). Bonferroni corrected (42 comparisons) P values <0.0012 shown in bold.

|  | STRING<br>Reference<br>(n=15561) | Born in Bradford<br>LOF all (n=476) | Decode<br>LOF all<br>(n=3769) | Decode<br>LOF “benign”<br>(n=3530) | Decode<br>LOF “pathogenic”<br>(n=416) | Orphanet<br>GOF all<br>(n=106) | Orphanet<br>LOF all<br>(n=458) |
| --- | --- | --- | --- | --- | --- | --- | --- |
| Binding<br>p-value, median,<br>[5<sup>th</sup>-95<sup>th</sup> quantiles] | 10, [0-139] | **9.3 x10<sup>-11</sup>**<br>**5.5, [0-61]** | **6 x10<sup>-07</sup>**<br>**9, [0-115]** | **2 x10<sup>-06</sup>**<br>**9, [0-106]** | 0.0105<br>7, [0-271] | **2.8 x10<sup>-09</sup>**<br>**26, [2-152]** | 0.0018<br>14, [0-95] |
| Reaction<br>p-value, median,<br>[5<sup>th</sup>-95<sup>th</sup> quantiles] | 0, [0-77] | 0.0047<br>0, [0-39] | 0.0152<br>0, [0-71] | 0.0019<br>0, [0-55] | 0.0134<br>0, [0-543] | **3 x10<sup>-06</sup>**<br>**0, [0-52]** | 0.0499<br>0, [0-73] |
| Activation<br>p-value, median,<br>[5<sup>th</sup>-95<sup>th</sup> quantiles] | 0, [0-6] | 0.533<br>0, [0-4] | **3.9 x10<sup>-06</sup>**<br>**0, [0-4]** | **1.9 x10<sup>-06</sup>**<br>**0, [0-4]** | 0.3139<br>0, [0-4] | **1 x10<sup>-17</sup>**<br>**1, [0-44]** | **3.3 x10<sup>-13</sup>**<br>**0, [0-11]** |
| Expression<br>p-value, median,<br>[5<sup>th</sup>-95<sup>th</sup> quantiles] | 0, [0-8] | 0.3723<br>0, [0-6] | **3.6 x10<sup>-05</sup>**<br>**0, [0-5]** | **4 x10<sup>-05</sup>**<br>**0, [0-5]** | 0.0309<br>0, [0-4] | **1.7 x10<sup>-17</sup>**<br>**1, [0-45]** | **1.8 x10<sup>-18</sup>**<br>**0, [0-23]** |
| Catalysis<br>p-value, median,<br>[5<sup>th</sup>-95<sup>th</sup> quantiles] | 0, [0-9] | **0.0007**<br>**0, [0-1]** | **0.0001**<br>**0, [0-4]** | **0.0003**<br>**0, [0-4]** | 0.019<br>0, [0-1] | **7.9 x10<sup>-08</sup>**<br>**0, [0-27]** | 0.0001<br>0, [0-17] |
| Post Trans Mod<br>p-value, median,<br>[5<sup>th</sup>-95<sup>th</sup> quantiles] | 0, [0-6] | 0.002<br>0, [0-3] | 0.0051<br>0, [0-4] | 0.0119<br>0, [0-5] | 0.00135<br>0, [0-3] | **6.4 x10<sup>-27</sup>**<br>**1.5, [0-31]** | **8.2 x10<sup>-05</sup>**<br>**0, [0-9]** |
| All Interactions<br>p-value, median,<br>[5<sup>th</sup>-95<sup>th</sup> quantiles] | 14, [0-211] | **3.4 x10<sup>-09</sup>**<br>**7, [0-98]** | **4.8 x10<sup>-08</sup>**<br>**11, [0-166]** | **4.7 x10<sup>-08</sup>**<br>**11, [0-155]** | 0.0438<br>10, [0-815] | **2 x10<sup>-12</sup>**<br>**47.5, [4-305]** | **1.1 x10<sup>-06</sup>**<br>**24, [1-180]** |

**Table S5.**
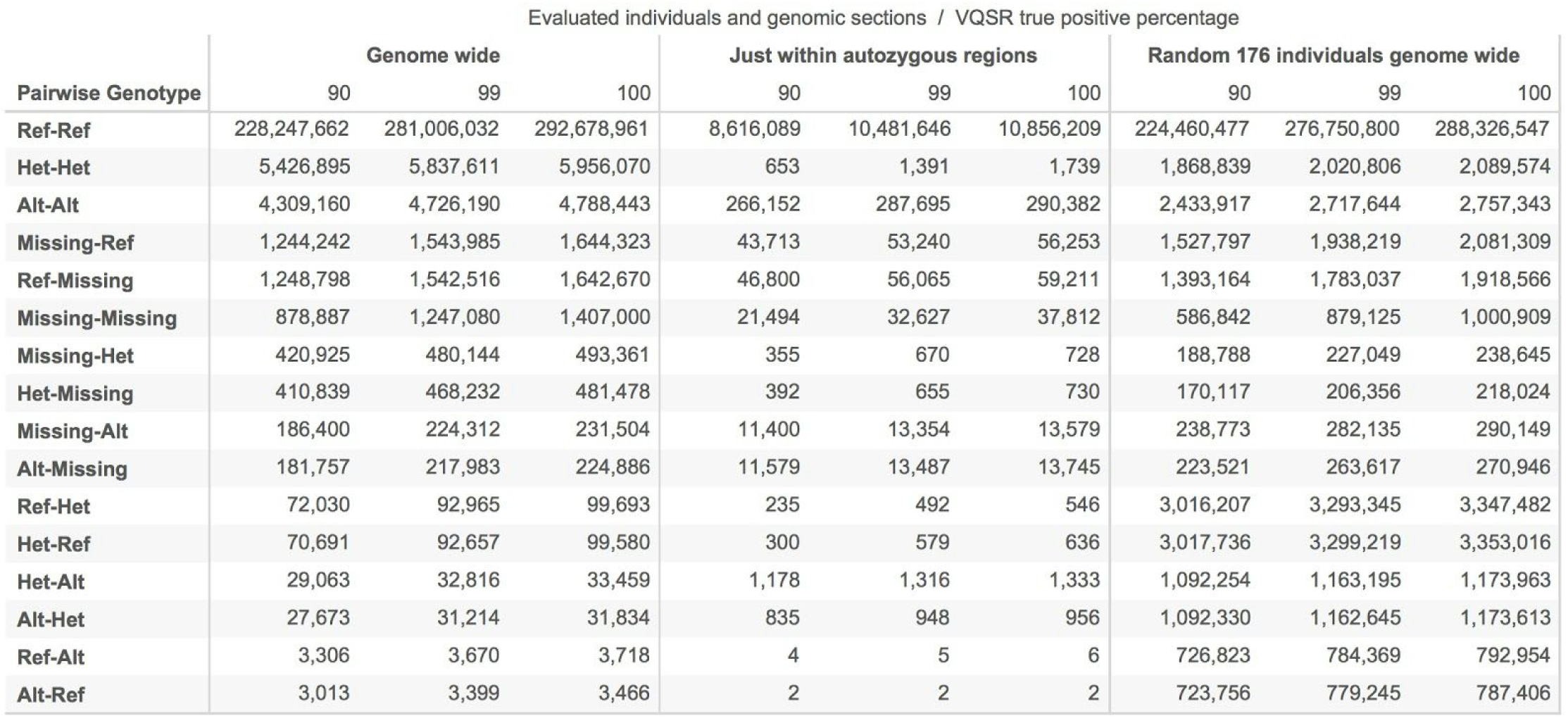
Pairwise genotypes for duplicate samples across evaluated individuals and regions. Pairwise genotypes (Ref, Het, Alt) across 176 duplicate pairs that were evaluated genome wide and just within the autozygous sections along with those from 176 unrelated (random) pairs. VQSR true positive percentage reflect the portion of true positive sites left (at VQSR= 90, 99, 100).

**Table S6.**
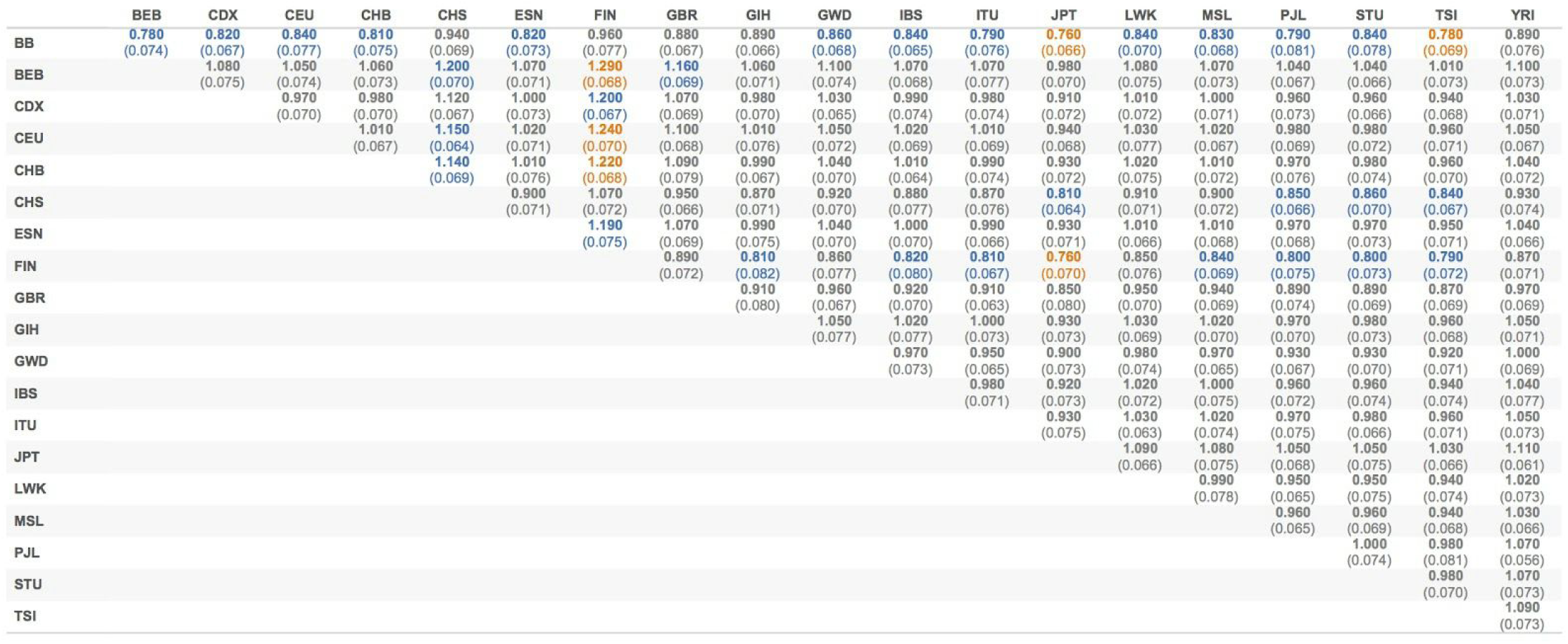
Pairwise R_A,B_ values across different populations. Numbers in bold reflect the pairwise R_A,B_ values for LOF variants normalized by the synonymous variant counts while the jacknife standard errors are shown in brackets. Highlighted in blue are comparisons that are at least 2 standard errors away from 1 (neutral expectation) and in orange are comparisons that are 3 standard errors away.

**Table S7.** 10XGenomics summary statistics on sequenced samples.

|  | SNPS phased | DNA in molecules<br>>20kb | GEMs detected | N50 phase block | Mean depth |
| --- | --- | --- | --- | --- | --- |
| NA12878 | 96.2% | 89.1% | 247924 | 16674432 bp | 33.9 X |
| NA12882 | 97.8% | 85.4% | 245880 | 9346942 bp | 28.2 X |
| PRDM9 knockout<br>mother | 92.6% | 70.4% | 235515 | 586148 bp | 29.9 X |
| her son | 90.7% | 27.9% | 223067 | 157393 bp | 26.2 X |

**Table S8.**
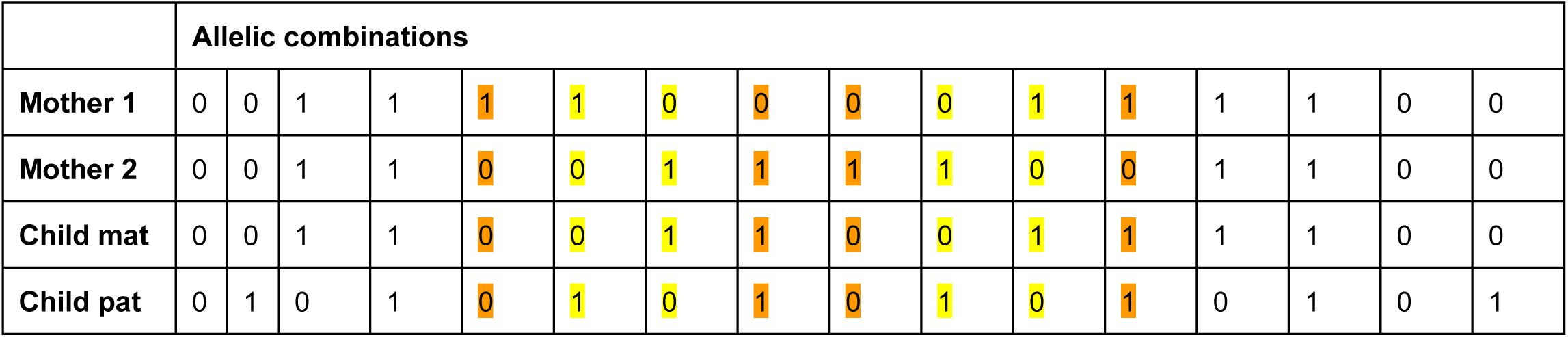
Allelic combinations of sites informative about recombination in mother child duos. Numbers show the different alleles (O-reference, 1-alternate) seen in the parent and child for bi-allelic sites. The first 8 configurations show transmission of the chromosome Mother 2 while the next 8 show transmission of chromosome Mother 1. Sites that are informative when only the mother is phased are shown in orange, while additional sites that are informative when the child is also phased so as to identify the maternal chromosome are shown in yellow.

